# Male fertility is independent of Enh13 control of *Sox9* testicular expression

**DOI:** 10.64898/2026.02.09.704748

**Authors:** Maor Lubman, Meshi Ridnik, Isabelle Stévant, Yael Kimchi Djanshvili, Elisheva Abberbock, Shelly Ziv Lhermann, Nitzan Gonen

## Abstract

Testis development relies on precise temporal control of *Sox9* expression, which is rapidly activated to initiate testis development and subsequently maintained to preserve Sertoli cell identity and male fertility. The distal enhancer, Enh13, is essential for *Sox9* upregulation as its constitutive deletion results in complete XY male-to-female sex reversal. Enh13’s requirement across distinct developmental stages remains unexplored. We show that early gonadal deletion of Enh13 fully recapitulates XY sex reversal, demonstrating that Enh13 activity is strictly required to initiate the male pathway. In contrast, Sertoli cell-specific deletion of Enh13 after sex determination has no effect on *Sox9* expression, testis architecture, or male fertility, revealing that Enh13 is dispensable for *Sox9* maintenance in the differentiated testis. Chromatin accessibility identifies candidate enhancers gaining activity after sex determination, suggesting a regulatory handoff within the *Sox9* locus. Finally, we show that duplicating Enh13 in the endogenous mouse locus fails to recapitulate the human XX sex reversal, suggesting that it may not be solely Enh13 sequence duplication that led to XX sex reversal in humans. To the best of our knowledge, this study represents the first *in vivo* conditional knockout of a developmental enhancer, allowing for temporally controlled interrogation of regulatory logic.

## Introduction

The mammalian testes represent a seminal component of the male reproductive system, responsible for storage of spermatogonial stem cells and generation of spermatozoa, along with production of androgens throughout embryonic and adult life (Wilhelm et al. 2007). The testes are composed of several cell types including somatic and germ cells, organised into seminiferous tubules, where sperm is produced, and an interstitial compartment. Among testicular somatic cells are Sertoli cells, which support and regulate the differentiation of germ cells within the seminiferous tubules, and Leydig cells, located in the interstitium, responsible for androgen production (Wilhelm et al. 2007). The *Sox9* gene, expressed in Sertoli cells, is considered a master regulator gene necessary for both testis development during embryonic stages, and later on, for proper testis function and male fertility (Barrionuevo et al. 2009; Lavery et al. 2011; Gonen and Lovell-Badge 2019).

Mammalian testes develop at embryonic stages from the genital ridge, a bipotential tissue that has the capacity to differentiate into either a testis or an ovary, depending on the genetic program activated at the time of sex determination (Wilhelm et al. 2007; Rotgers et al. 2018; Stevant and Nef 2019; Wilhelm et al. 2025). In XY embryos, the expression of the Y-encoded SRY transcription factor in supporting cell precursors initiates Sertoli cell differentiation and testis development by upregulating the expression of SRY’s direct target gene, *Sox9* (Sinclair et al. 1990; Koopman et al. 1991; Sekido et al. 2004). Once expressed above a certain threshold, SOX9 is necessary and sufficient to induce testis development (Gonen and Lovell-Badge 2019). Indeed, mice and human individuals carrying loss-of-function alleles of *Sox9/SOX9* present XY male-to-female sex reversal (Foster et al. 1994; Wagner et al. 1994; Chaboissier et al. 2004; Lavery et al. 2011), whereas overexpression of *Sox9/SOX9* in the context of XX gonads leads to the opposite, XX female-to-male sex reversal (Huang et al. 1999; Vidal et al. 2001).

The expression and function of *Sox9* in the murine gonads can be divided into three main stages; In the early bipotential gonads, at embryonic day (E)10.5, *Sox9* is expressed at low, basal levels, in both XY and XX gonads prior to sex determination (Morais da Silva et al. 1996; Zhao et al. 2018).

Next, at the stage of sex determination, at around E11.0-E11.5, *Sox9* expression is markedly upregulated in Sertoli cells due to joint activity of SRY and SF1, while being actively repressed in pre-granulosa cells of the ovary (Sekido et al. 2004; Zhao et al. 2018; Stevant et al. 2025). This strong *Sox9* activation is mandatory for testis development, while the repression is required for proper ovarian specification (Wilhelm et al. 2007; Rotgers et al. 2018; Stevant and Nef 2019; Wilhelm et al. 2025). As such, early conditional removal of *Sox9* in the gonads using the *Sf1*-Cre line led to complete XY male-to-female sex reversal (Chaboissier et al. 2004; Lavery et al. 2011). Once expressed above a certain threshold, SOX9 is involved in *Sry* repression as well as auto-activating its own expression together with SF1 (Sekido and Lovell-Badge 2009; Gonen and Lovell-Badge 2019). Additional signalling pathways are involved in maintaining high *Sox9* levels including the FGF9 (Schmahl et al. 2004; Kim et al. 2006; Kim et al. 2007), PGD2 (Wilhelm et al. 2005) and the partly redundant SOX8 transcription factor (Barrionuevo et al. 2009).

Unlike *Sry*, which is only transiently expressed in the testis at the stage of sex determination, *Sox9* expression persist in the testis throughout life and is fully required for Sertoli cell maintenance and fertility. Conditional removal of *Sox9* after the stage of sex determination, using the *Amh*-Cre line (E14.0), resulted in mice with normal testis development which were initially fertile however, similarly to the *Sox8* null mice (Sock et al. 2001; O’Bryan et al. 2008), became sterile at around 5 months of age (Barrionuevo et al. 2009). Breeding the late gonadal *Sox9* conditional knockout (cKO) on a *Sox8* null allele led to complete primary infertility, highlighting a partial redundancy between SOX9 and SOX8 in Sertoli cell fate maintenance and testis function (Barrionuevo et al. 2009). It also showed that SOX9/SOX8 are fully necessary for proper testicular function.

As SOX9 is an important transcription factor, needed for the development and function of many tissues including cartilage, brain, pituitary, lung, heart, testis and others, much interest was drawn to the regulatory elements involved in its spatiotemporal gene expression (Gonen and Lovell-Badge 2019). Several enhancers involved in testicular *Sox9* expression during sex determination have been identified including TES, Enh8, Enh13, Enh14 and Enh32, all of which are located in the 2 MB gene desert upstream of the *Sox9* gene (Sekido and Lovell-Badge 2008; Gonen et al. 2018; Gonen and Lovell-Badge 2019). While many of these enhancers seem to function in a redundant manner (Gonen et al. 2017; Gonen et al. 2018), Enh13 has proven to be crucial for *Sox9* induction in the Sertoli cells. XY mice carrying deletion of Enh13, or small mutations in the SOX9 and SRY or GATA4 transcription factor binding sites (TFBS) of Enh13, present with complete XY female sex reversal, fully phenocopying mutations in the *Sox9* gene itself (Gonen et al. 2018; Ogawa et al. 2018; Ridnik et al. 2024; Ogawa et al. 2025). Enh13 was shown to be the docking site for SRY/ SOX9/ SF1 binding in a way that strongly upregulates *Sox9* expression (Gonen et al. 2018; Ridnik et al. 2024). Human individuals carrying deletion of the XYSR region, in which Enh13 is fully embedded within, also present with XY female development, suggesting that Enh13 function in mediating *Sox9* expression during sex determination is highly conserved in mammals (Baetens et al. 2017; Croft et al. 2018; Gonen et al. 2018; Ogawa et al. 2018; Ridnik et al. 2024). While many gonadal enhancers of *Sox9* have been described to date, we still do not know what regulates the early gonadal basal level of *Sox9,* nor do we know what regulates testicular *Sox9* expression maintenance after sex determination.

Enhancers are considered to regulate their target genes in a temporal and spatial manner. It is suggested that a single gene can have different enhancers in different tissues, and even different enhancers across distinct stages of development of the same tissue (Long et al. 2016; Spitz 2016; Schoenfelder and Fraser 2019; Bolt and Duboule 2020; Ibrahim and Mundlos 2020). While Enh13 has been shown to be critical for proper *Sox9* expression during the sex determination stage, it remains unknown if this potent enhancer continues to be critical also for later stages of *Sox9* gene expression maintenance or if other regulatory elements take over that role.

Several XX human individuals with Differences of Sex Development (DSD) that carry small duplication encompassing the human homologue of Enh13 (eSR-A) were described as ovotesticular/ testicular DSD (Croft et al. 2018; Sajan et al. 2023). While the mechanism of action was never experimentally resolved *in vivo*, it was speculated that in the presence of two copies of Enh13/ eSR-A, the basal levels of SOX9, normally present at the early bipotential stage, can enhance *SOX9* expression beyond the threshold level that would lead to testis differentiation, in the absence of SRY (Gonen and Lovell-Badge 2019; Sreenivasan et al. 2022). It remains unknown if this hypothesis is correct and whether duplication of a regulatory element can dramatically induce gene expression of the target gene.

Here, we generated several genome-edited mice models to explore the late gonadal role of Enh13, as well as copy number variation. We first generated a conditional floxed allele of Enh13. This allele was bred with two Cre lines: one that induced Enh13 deletion in the testis during sex determination, and one that induced deletion after sex determination. Early conditional loss of Enh13 led to fully penetrant XY male-to-female sex reversal, similar to full Enh13 knock out. In contrast, late loss of Enh13 in Sertoli cells resulted in XY fertile mice with normal testis and no change in *Sox9* expression levels. This suggests that while Enh13 is an extremely critical enhancer of *Sox9* at the time of sex determination, it is dispensable for the maintenance of *Sox9* expression and male fertility. Exploring our recent ATAC-seq data, along with published ATAC-seq datasets, we could identify novel potential *Sox9* enhancers that may function in the maintenance of *Sox9* in the testis to allow proper testicular function and fertility. Next, to model the 46, XX DSD patients carrying duplication of Enh13, we generated mice carrying two tandem repeats of Enh13 in the endogenous genomic locus. XX and XY mice presented with normal ovary and testis development at embryonic and adult stages, respectively. This suggest that simple duplication of Enh13 is insufficient to explain human 46, XX DSDs associated with duplication of the Enh13-containing region.

## Results

### Early conditional loss of Enh13 in the gonads results in XY sex reversal

A full KO of Enh13 results in XY male-to-female sex reversal in mice and humans (Croft et al. 2018; Gonen et al. 2018). To explore if also early gonad conditional KO of Enh13, solely in somatic gonadal cells, would result is the same fully-penetrant XY sex reversal, we generated a floxed allele of Enh13 (Supplemental Fig. S1A). We performed a two-step homology dependent repair (HDR) CRISPR electroporation in zygotes, first inserting the 5’ LoxP site, followed by the 3’ LoxP site. Wild type (WT) C57BL6 zygotes were electroporated with a single guide RNA (sgRNA) targeting the 5’ side outside Enh13 along with a single strand oligo (ssOligo) containing a LoxP site (34 bp) and two short homology arms. Founder mice carrying the 5’ LoxP site were bred to WT mice for germline transmission, and two heterozygous mice were bred to homozygosity. Next, sperm was retrieved from males homozygous for the 5’ LoxP insertion and was used to fertilize WT oocytes using In Vitro Fertilization (IVF), making all resulting zygotes heterozygous for the 5’ LoxP site. These IVF-generated zygotes were subjected to CRISPR electroporation with a sgRNA targeting the 3’ side of Enh13 along with a ssOligo containing a LoxP site and two short homology arms. Founder mice from this step were screened for the presence of two LoxP sites on the same allele, and positive founders were bred to WT mice for germline transmission. Two heterozygous mice were bred to homozygosity and the presence of the two LoxP sites spanning the endogenous Enh13 was validated using long PCR Sanger sequencing (Supplemental Fig. S1B, Materials and Methods).

Next, these floxed Enh13 mice were bred to the *Sf1*-Cre mouse line that induces early removal of Enh13 in somatic cells of the gonad (Bingham et al. 2006) (Fig. 1A). Two control mice were generated: mice homozygous for the floxed Enh13 without the Cre (Control 1, Enh13^fl/fl^) and mice carrying the *Sf1*-Cre and only one allele of floxed out Enh13 (Control 2, Enh13^fl/+^; *Sf1*-Cre^+^). Experimental mice carried the *Sf1*-Cre and two alleles of floxed out Enh13, fully removing Enh13 presence from early somatic gonadal cells (Experimental, Enh13^fl/fl^; *Sf1*-Cre^+^) (Fig. 1A). Analysis of these mice at adulthood revealed that the two control XY mice presented as males externally and internally, whereas mice with early gonadal conditional removal of Enh13 exhibited fully penetrant XY male-to-female sex reversal (N=10) (Fig. 1B). These mice presented with female external genitalia, marked by short anogenital distance and presence of nipples, similar to XX WT female. Internally, these XY mice exhibited ovaries, oviduct and uterus, indistinguishable from that of WT XX mice (Fig. 1B). Analysing adult gonads further using immunostaining confirmed complete sex reversal with XY Enh13^fl/fl^; *Sf1*-Cre^+^ gonads presenting FOXL2 expression in granulosa cells, lacking SOX9 expression, normally present in Sertoli cells, and displaying follicular structure containing TRA98-positive germ cells (Fig. 1C).

**Figure 1.**
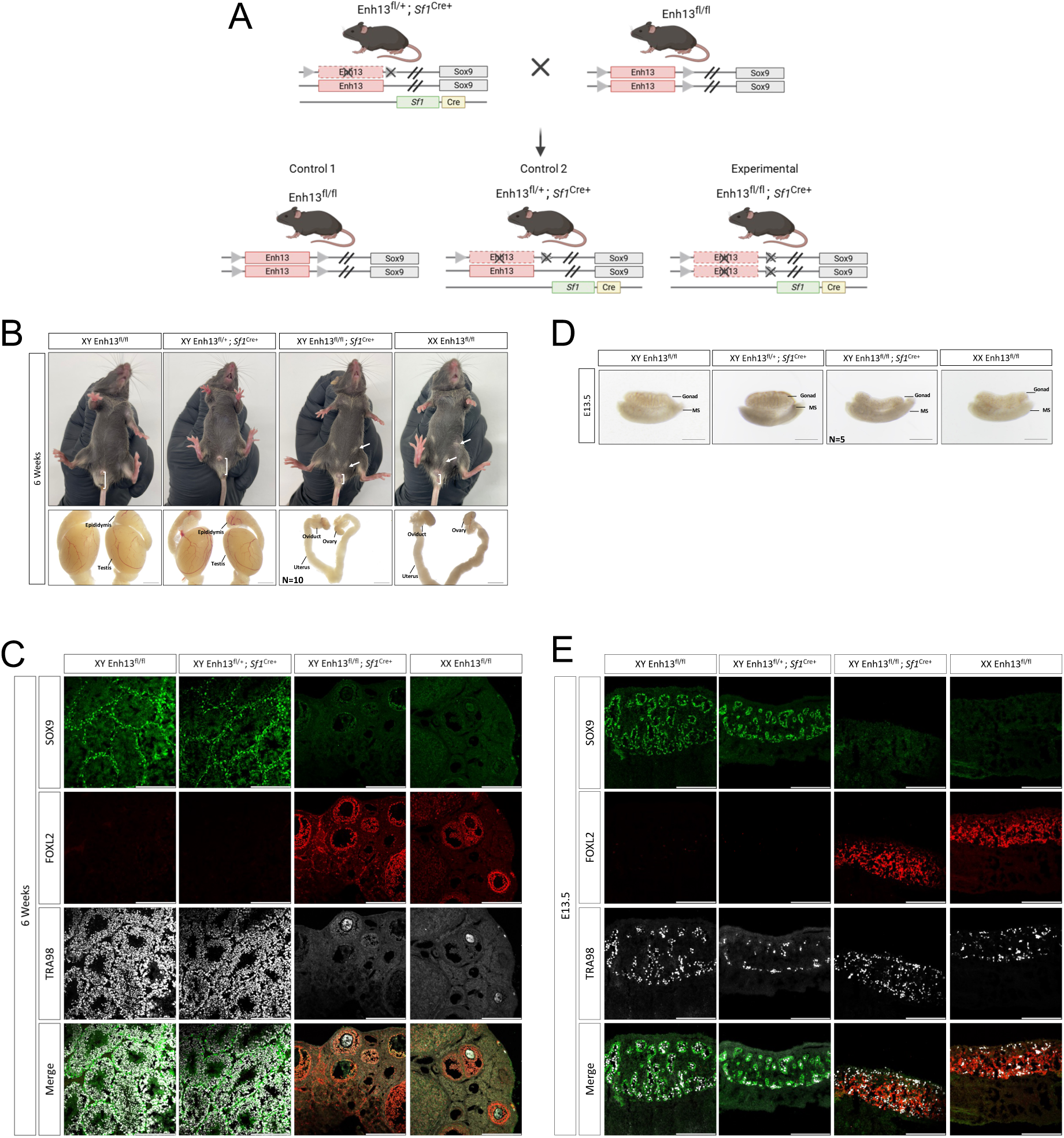
Early loss on Enh13 in the gonads leads to XY sex reversal. (A) Schematic representation of breeding scheme set up to generate experimental and control mice. (B) Bright field images of the external and internal genitalia and gonads of 6-Week-old adult control mice (XY Enh13^fl/fl^, XY Enh13^fl/+^; *Sf1*^Cre+^, and XX Enh13^fl/fl^) and experimental mice (XY Enh13^fl/fl^; *Sf1*^Cre+^). Scale bar represents 2000 µm (C, E) Immunostaining of 6-Week-old gonads (C) or E13.5 gonads (E) from control and experimental mice. Gonads were stained for Sertoli-marker SOX9 (green), granulosa-marker FOXL2 (red) and germ cell marker TRA98 (grey). Scale bars represent 200 µm. (D) Bright field images of E13.5 gonads of control and experimental mice. Scale Bar represents 500 µm.

Next, we wanted to explore if this sex reversal, present in adult mice, also appear as fully penetrant XY sex reversal already from embryonic stages. To that aim we dissected E13.5 gonads from the two XY controls, XY experimental and XX control embryos. While both XY control embryos presented an embryonic testis with testis cords and a coelomic vessel, all XY Enh13^fl/fl^; *Sf1*-Cre^+^ embryos analysed (N=5) presented a gonad representing an ovary, devoid of testis cords and a coelomic vessel, similarly to XX control ovary (Fig. 1D). To explore if this sex reversed gonad was an ovary or an ovotestis, we performed immunostaining with SOX9, FOXL2 and TRA98. While both XY control gonads displayed SOX9 expressing Sertoli cells, organised into testis cords, engulfing TRA98-positive germ cells, XY Enh13^fl/fl^; *Sf1*-Cre^+^ gonads failed to express SOX9 and instead expressed FOXL2 and TRA98, in a classical ovarian presentation, as in XX control gonad (Fig. 1E).

Altogether, these results suggest that early conditional KO of Enh13, solely in the somatic cells of the gonad, is able to fully recapitulate the complete XY male-to-female sex reversal observed upon full KO of Enh13 (Gonen et al. 2018). This also confirms the effectiveness of the newly generated floxed Enh13 allele.

### Late conditional loss of Enh13 in the gonads does not lead to male infertility

While SOX9 is crucial to allow testis sex determination, it is also mandatory later on, along with SOX8, for the maintenance of Sertoli cell fate throughout life, and hence proper male fertility (Barrionuevo et al. 2009; Gonen and Lovell-Badge 2019). We therefore wanted to explore whether Enh13, which has proven to be the sole critical enhancer controlling *Sox9* expression at the stage of sex determination, continues to play a critical role in regulating *Sox9* expression beyond the stage of sex determination. To address this, we bred the Enh13 floxed allele with the *Amh*-Cre allele that induced Cre activity specifically in Sertoli cells, from E14.0 (Fig. 2A) (Lecureuil et al. 2002). This same Cre driver was previously used to assess the late functions of *Sox9* in Sertoli cell fate maintenance and male fertility (Barrionuevo et al. 2009). Similarly to above, this breeding generated two control mice: mice carrying two floxed Enh13 alleles without the Cre (Control 1, Enh13^fl/fl^) and mice carrying the *Amh*-Cre and one allele of floxed out Enh13 (Control 2, Enh13^fl/+^; *Amh*-Cre^+^). Experimental mice carried two floxed out alleles of Enh13 along with the *Amh*-Cre, removing Enh13 in Sertoli cells from E14.0 (Experimental, Enh13^fl/fl^; *Amh*-Cre^+^) (Fig. 2A). Adult mice were analysed at two stages: 6 weeks and 6 months. This is due to the fact that late conditional removal of *Sox9* itself (using *Amh*-Cre) resulted in XY mice with small testes starting from 5 months of age. Analysis of adult XY males from the two controls and experimental mice demonstrated that both the XY controls and experimental males presented with long anogenital distance and male external and internal appearance at both 6 weeks (Fig. 2B) and 6 months (Fig. 2C). At 6 weeks and 6 months, experimental males exhibited testis at similar size and weight to the control males (Fig. 2B-D). We next wanted to assess the state of the adult gonads and performed immunostaining of 6 weeks (Fig. 2E) and 6 months (Fig. 2F) gonads. Immunostaining revealed SOX9 expression in Sertoli cells surrounding the seminiferous tubules, DDX4-positive germ cells were present in the testis and SF1-positive Leydig cells were positioned in the interstitial compartment.

**Figure 2.**
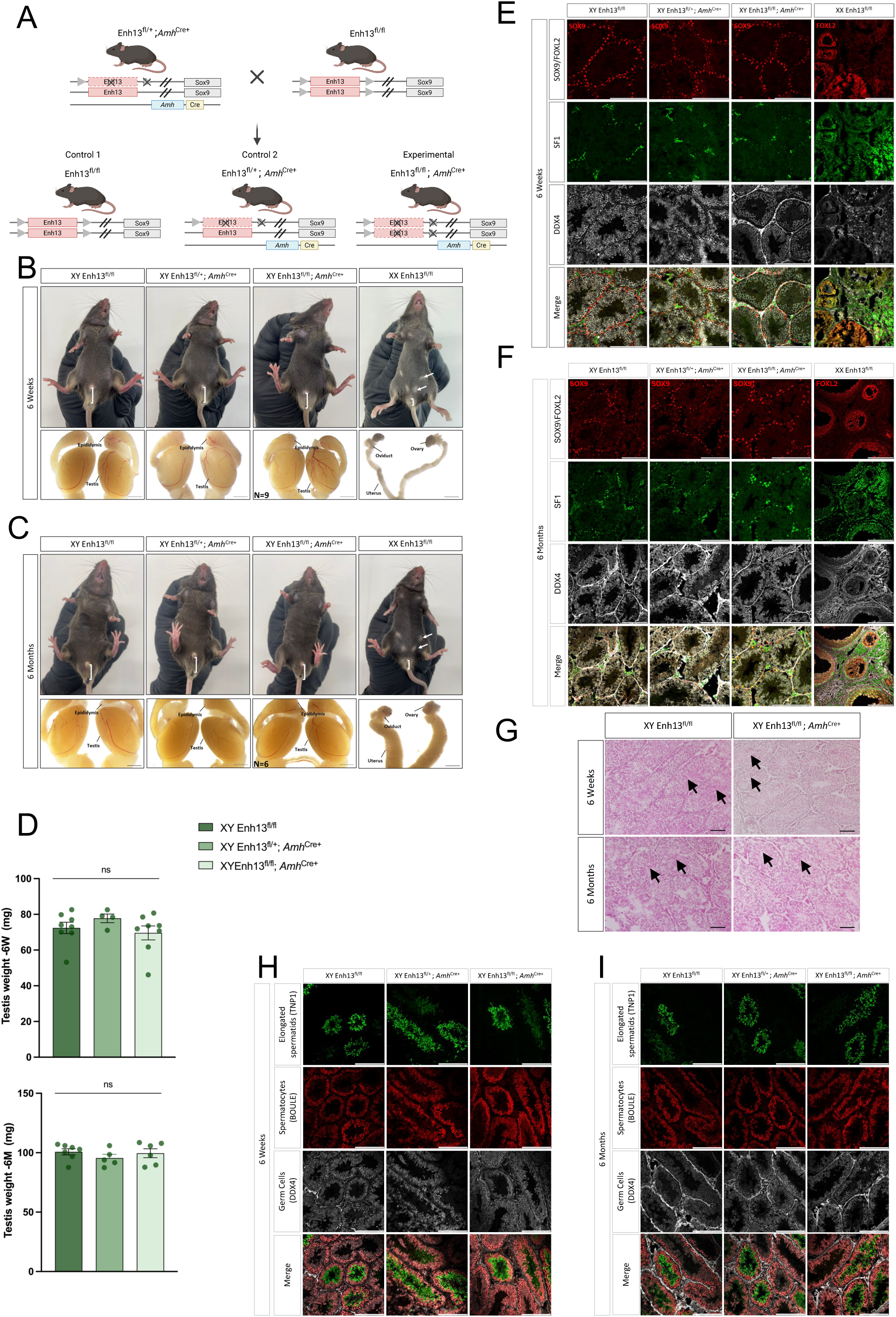
Late loss on Enh13 in the Sertoli cells does not result in male infertility. (A) Schematic representation of breeding scheme set up to generate experimental and control mice. (B-C) Bright field images of the external and internal genitalia and gonads of 6-Week-old (B) and 6-Month-old (C) adult control mice (XY Enh13^fl/fl^, XY Enh13^fl/+^; *Amh*^Cre+^, and XX Enh13^fl/fl^) and experimental mice (XY Enh13^fl/fl^; *Amh*^Cre+^). Scale bar represents 2000 µm. (D) Quantification of testis weight at 6-Week-old and 6-Month-old mice. Each point represents the average weight of both testes of an individual mouse; error bars show SEM. Statistical analysis was performed using one-way ANOVA tests in Prism 10 software. P values below 0.05 were considered statistically significant. ns- not significant (E-F,H-I) Immunostaining of 6-Weeks-old gonads (E,H) and 6-Month-old gonads (F,I) from control and experimental mice. Scale bars represent 200 µm. (E-F) Gonads were stained for either Sertoli-marker SOX9 (testis) or granulosa-marker FOXL2 (ovaries) (red), Leydig/ theca cell marker SF1 (green) and germ cells marker DDX4 (grey). (H-I) Testis were stained for elongated spermatid marker TNP-1 (green), spermatocyte marker BOULE (red), and germ cells marker DDX4 (grey). (G) H&E-stained sections of 6-Week-old and 6-Month-old old testis of control and experimental mice. Arrows indicate seminiferous tubules containing elongated spermatids. Scale bars represent 100 µm.

To examine if these mice have elongated spermatids, we performed H&E histology staining on 6 weeks and 6 months old testis sections (Fig. 2G). This analysis revealed that experimental mice had elongated spermatids within the seminiferous tubules. To validate this, we also performed immunostaining of 6 weeks and 6 months old testis with Transition Protein 1 (TNP1), which marks elongated spermatids, BOULE, which marks spermatocytes and DDX4 (Fig. 2H-I). Immunostaining confirmed the presence of spermatocytes and elongated spermatids at both 6 weeks (Fig. 2H) and 6 months old testis (Fig. 2I) and indeed these mice were able to give rise to pups, establishing their normal fertility.

Analysis of XX mice carrying the floxed allele of Enh13 with *Amh*-Cre did not reveal any observed phenotype, although *Amh*-Cre is active in adult ovaries, again, supporting that Enh13 has no function in adult gonads (Supplemental Fig. 2) (Lecureuil et al. 2002).

Overall, these results suggest that while Enh13 play a critical role at regulating *Sox9* expression during the sex determination stage, it is redundant or not functional during the maintenance of *Sox9* expression in the testis after sex determination.

### Enh13 is dispensable for testicular Sox9 expression beyond sex determination and other enhancers may take over this role

To assess more directly if *Sox9* expression is altered due to removal of Enh13 after the sex determination stage we performed Real Time quantitative PCR on gonads of 6 weeks old XY controls and experimental mice, along with XX control mice. No significant change in the expression levels of *Sox9*, and its partner, *Sox8,* could be seen between XY Enh13^fl/fl^, Enh13^fl/+^; *Amh*-Cre^+^ and Enh13^fl/fl^; *Amh*-Cre^+^ gonads (Fig. 3A). This indicates that Enh13 is dispensable for the maintenance of *Sox9* expression after sex determination.

**Figure 3.**
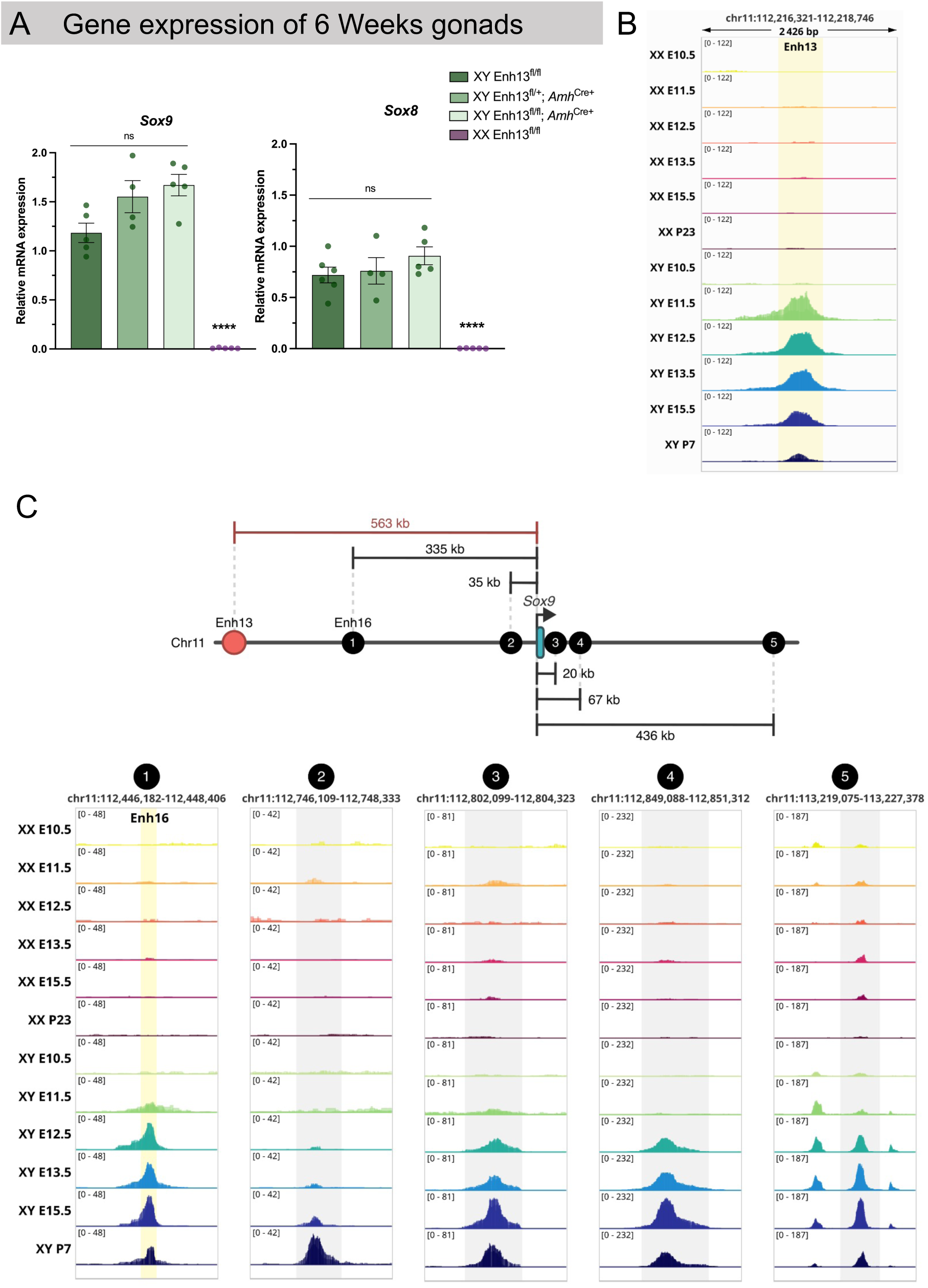
Enh13 does not control *Sox9* expression post sex determination and may be replaced by other putative enhancers. (A) Real-time quantitative PCR analysis (qRT-PCR) of genes involved in male Sertoli cells maintenance (*Sox9* and *Sox8*) in gonads from 6 Weeks-old mice of control (XY Enh13^fl/fl^, XY Enh13^fl/+^; *Amh*^Cre+^, and XX Enh13^fl/fl^) and experimental mice (XY Enh13^fl/fl^; *Amh*^Cre+^). Data are presented as mean 2^-ΔΔCt^ values ±SEM, normalized to the housekeeping gene *Hprt*. Statistical analysis was done using one-way ANOVA followed by Dennett’s posttest. Samples were compared to XY Enh13^fl/fl^. *P < 0.05, **P < 0.01, ***P < 0.001, and ****P < 0.0001, ns-not significant. (B) ATAC-seq profiles at the Enh13 locus (chr11:112,216,321–112,218,746) in XX and XY gonads somatic cells of E10.5 as well as purified Sertoli and pre-granulosa cells from E11.5 to postnatal stages. Enh13 (yellow) shows strong, transient accessibility in XY gonads during the sex-determining window (E11.5–E13.5), with decreased accessibility at P7. There is little or no Enh13 accessibility in XX gonads. (C) Genomic organization of putative regulatory regions surrounding *Sox9*, including Enh13, Enh16, and additional distal elements (regions 1–5), with inter-element distances indicated. ATAC-seq signal at each putative enhancer across XX and XY gonads somatic cells of E10.5 as well as purified Sertoli and pre-granulosa cells from E11.5 to postnatal stages reveals distinct temporal and sex-specific accessibility patterns, highlighting a complex regulatory landscape upstream of *Sox9*.

To better explore *Sox9* regulatory landscape before and after sex determination, we integrated previously published ATAC-seq datasets performed on somatic cells of the gonad, notably Sertoli and pre-granulosa cells. This included E10.5 XX and XY somatic gonadal cells (Gonen et al. 2018; Garcia-Moreno et al. 2019), E11.5 – E15.5 purified Sertoli and pre-granulosa cells (Stevant et al. 2025) and P7 enriched Sertoli cells as well as P23 enriched granulosa cells (Lindeman et al. 2021). Examining the genomic region of Enh13, it is not accessible at E10.5 in XY somatic cells, gaining strong accessibility in XY E11.5 – E13.5 Sertoli cells, while exhibiting decreased accessibility in E15.5 and more so, in P7 Sertoli cells (Fig. 3B). Enh13 is not accessible in pre-granulosa and granulosa cells throughout development (Fig. 3B). These findings support our results that Enh13 is mostly active at the sex determination step and is dispensable afterwards. Next, relying on the ATAC-seq data, we interrogated the upstream and downstream genomic region surrounding the *Sox9* gene to identify candidate enhancers that may gain accessibility after sex determination (E11.5) and maintain accessibility in late embryonic stages and pre-pubertal. We could identify five putative enhancers, two located upstream of *Sox9* and three located downstream of *Sox9* that present increased accessibility at E15.5/ P7 purified Sertoli cells (Fig. 3C). The first upstream enhancer, previously termed Enh16 (Gonen et al. 2018) (marked as 1 here) is located 335 kb upstream of *Sox9*. This enhancer gains accessibility at E12.5 in Sertoli cells, and is strongly accessible at E15.5, and to lesser extent at P7. Another candidate enhancer (marked as 2), located 35 kb upstream of *Sox9,* is only starting to gain accessibility at E15.5 with much more accessibility at P7 Sertoli cells. Looking downstream to *Sox9*, we identified three enhancers located 20 kb, 67 kb and 436 kb downstream (marked as 3, 4 and 5, respectively) which are not accessible at E10.5 and E11.5 XY cells but acquire accessibility from E12.5 onwards with the strongest peaks appearing at E15.5 and P7 Sertoli cells.

It is possible that one of these enhancers, some of them, or all of them together, cooperate to maintain the high *Sox9* expression levels needed in Sertoli cells after sex determination in order to maintain Sertoli cell identity and hence proper testicular function, spermatogenesis support and male fertility.

### Different outcome upon Enh13 duplication between mouse and human

After seeing that Enh13 only functions to regulate *Sox9* expression during sex determination, we sought to explore another open question regarding Enh13, which is the outcome of altering Enh13 copy number. Three 46, XX DSD (testicular and ovo-testicular) individuals were described that carry a small duplication upstream of the human *SOX9* gene, fully encompassing the human homologue of Enh13/eSR-A (Croft et al. 2018; Sajan et al. 2023). While it is clear why loss of Enh13 will prevent *Sox9/SOX9* expression and testis development, being the docking site for SRY and later on, SOX9 binding, it remains unclear why having two copies of the enhancer could induce testis development in the context of XX gonads. It was speculated that in the presence of two copies of Enh13, the basal level of SOX9, present at XX and XY early gonads, could enhance *SOX9* expression beyond the threshold needed to initiate testis development (Sreenivasan et al. 2022). To experimentally model this *in vivo* and also evaluate whether XX mice carrying duplicated Enh13 will present similar phenotype to that of human XX female-to-male sex reversal, we sought to generate an allele with two tandem copies of Enh13 in the endogenous *Sox9* locus of the mouse.

Mice carrying duplicated copy of Enh13 were generated using the CRIPSR-READI technique (Chen et al. 2019). C57BL6 zygotes were pre-incubated with Adeno Associated Viruses 6 (AAV6) carrying two copies of Enh13 along with 400 bp homology arms on either side (Supplemental Fig. 3A). Following incubation, zygotes were electroporated with two guides located outside of Enh13 to remove the endogenous Enh13 and replace it with the HDR template containing instead two Enh13 sequences in tandem. Founder mice carrying the duplicated Enh13 were genotyped and validated for the presence of the duplicated allele using long range PCR reaction followed by Sanger sequencing (Supplemental Fig. 3B). Positive founder mice were bred to WT mice to allow germline transmission and two heterozygous were bred to homozygosity. XX adult mice carrying a heterozygous or homozygous duplication of Enh13 presented as females with short anu-genital distance and nipples externally, and a feminized internal reproductive system including ovaries, oviduct and uterus (Fig. 4A). No change in ovary size or weight was evident between XX WT, heterozygous and homozygous for the duplicated Enh13 (Fig. 4A-B). We next performed immunostaining on adult, 6 weeks old, gonads and could observe ovaries expressing FOXL2 within granulosa cells, encompassing TRA98-positive oocytes. No sign for SOX9 expression was evident (Fig. 4C).

**Figure 4.**
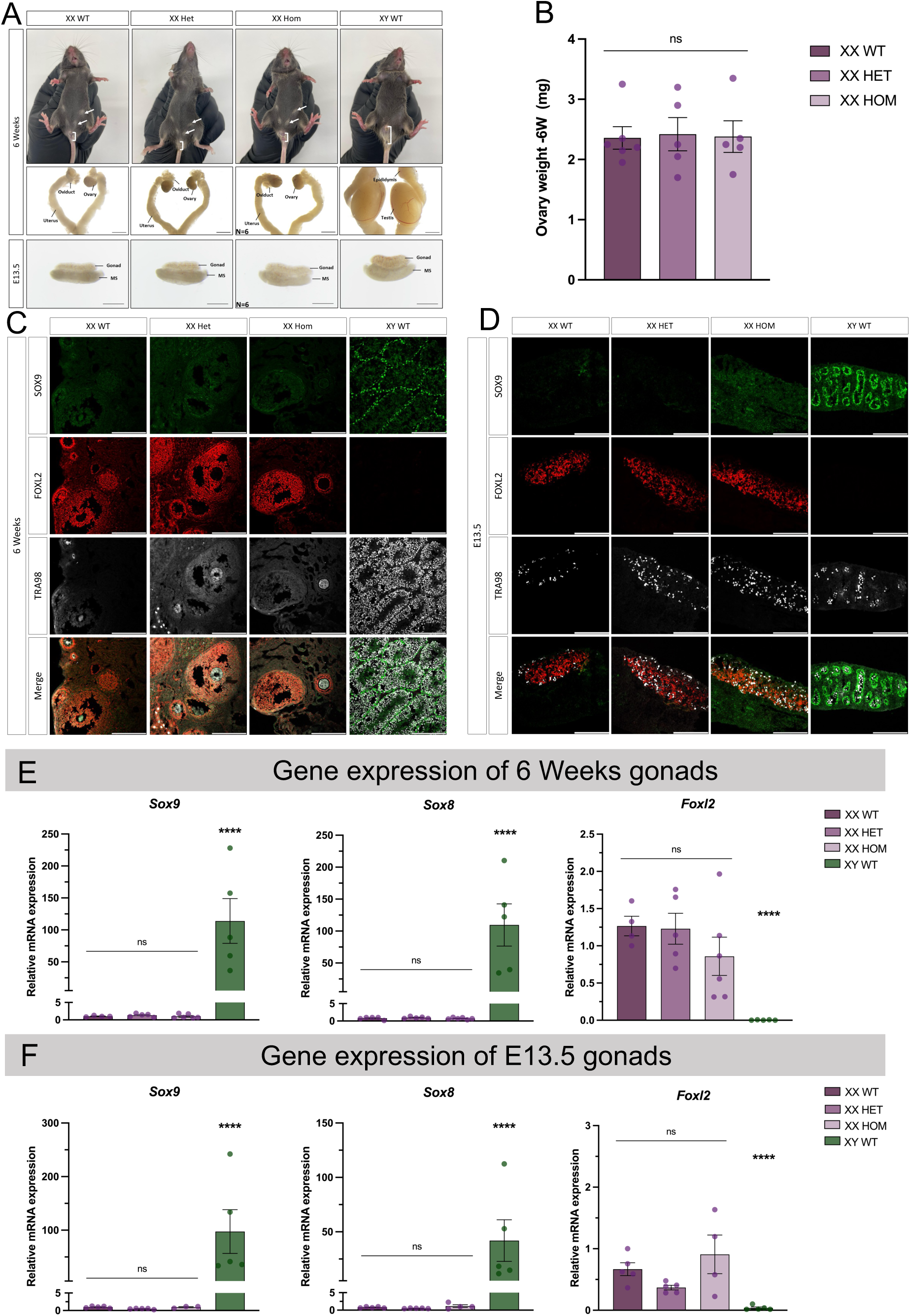
Duplication of Enh13 in mice does not phenocopy the phenotype of 46, XX DSD individuals. (A) Bright field images of the external and internal genitalia and gonads of 6-Week-old adult and E13.5 embryos of XX WT, XY WT, and XX heterozygous and homozygous mice for Enh13 duplication. Scale Bar represents 2000 µm in adults and 500 µm in embryonic gonads. (B) Quantification of ovary weight in 6-Week-old mice. Each dot represents the average weight of both ovaries of an individual mouse; error bars show SEM. Statistical analysis was performed using one-way ANOVA test in Prism 10 software where samples were compares to XX WT. P values below 0.05 were considered statistically significant. ns- not significant. (C-D) Immunostaining of 6-Week-old gonads (C) and E13.5 gonads (D). Gonads were stained for the Sertoli-marker SOX9 (green), granulosa-marker FOXL2 (red) and germ cells marker TRA98 (grey). Scale bars represent 200 μm. (E-F) Real-time quantitative PCR analysis of genes involved male (*Sox9* and *Sox8*) and in female (*Foxl2*) gonadal sex determination in gonads from 6-Week- old mice (E) and E13.5 embryos (F). Data are presented as mean 2^-ΔΔCt^ values, normalized to the housekeeping gene *Hprt*. Sample size of each genotype is indicated as dots representing the number of individuals. Error bars show SEM of 2^-ΔΔCt^ values. Statistical analysis was done by one-way ANOVA followed by Dennett’s posttest. \**P* < 0.05, \*\**P* < 0.01, \*\*\**P* < 0.001, and \*\*\*\**P* < 0.0001, ns- not significant. WT- *Wild Type*.

Since no phenotype appeared in XX adult gonads, we next examined E13.5 embryonic gonads (Fig. 4A). XX homozygous embryonic gonads were indistinguishable from XX WT or heterozygous gonads and presented with ovarian morphology, devoid of testis cords and a coelomic vessel (Fig. 4A). Immunostaining of embryonic gonads confirmed they express FOXL2 in pre-granulosa cells as well as TRA98 in germ cells. No sign for SOX9 expression was evident, ruling out the presence of ovotestis (Fig. 4D). Although no XX sex reversal was present at either adult or embryonic stages, we wanted to assess whether having two copies of Enh13 in XX gonads could increase *Sox9* expression. To that aim we performed RT-PCR on 6 weeks (Fig. 4E) or E13.5 embryonic gonads (Fig. 4F) of XX WT, XX het, XX hom and XY WT embryos. Analysis of gene expression revealed no elevation in the levels of either *Sox9*, or its direct target gene, *Sox8*. We could also not observe any major decrease in the expression levels of *Foxl2* (Fig. 4E-F).

While no elevation in *Sox9* expression levels were present in XX gonads, we were wondering whether duplicated Enh13 can lead to change in XY homozygous gonads. Analysis of adult XY mice carrying one or two alleles of duplicated Enh13 demonstrated normal external and internal male appearance (Supplemental Fig. 4A). Adult testis weight was identical to WT testis (Supplemental Fig. 4B). Embryonic gonads also appeared similar to XY WT gonads displaying testis cords and coelomic vessel.

Altogether, this analysis reveals that in mice, having two copies of Enh13 in the endogenous *Sox9* locus does not lead to increased *Sox9* expression and XX female-to-male sex reversal as seen in 46, XX DSD individuals. This suggests either a different sensitivity to Enh13 duplication between mice and human or that the sex reversal phenotype seen in human 46, XX DSD individuals does not stem solely from Enh13 copy number changes.

## Discussion

The *Sox9/SOX9* gene is a highly regulated transcription factor, expressed, and often serves as a master regulator, in many different tissues during development and disease (Gonen and Lovell-Badge 2019; Sreenivasan et al. 2022). Many pathological conditions have been described in relation to modifications of the *SOX9* regulatory landscape in various tissues including cartilage, craniofacial tissues, hair follicle, and the gonads (Franke et al. 2016; Baetens et al. 2017; Liu et al. 2017; Gonen and Lovell-Badge 2019; Long et al. 2020; Yang et al. 2023). We have previously extensively investigated the regulatory elements involved in the testicular expression of *Sox9* (Sekido and Lovell-Badge 2008; Gonen et al. 2017; Gonen et al. 2018; Ridnik et al. 2024) and identified Enh13 as a critical gonadal enhancer of *Sox9*, fully needed for testis development (Gonen et al. 2018; Ogawa et al. 2018). In this study, we show that early conditional deletion of Enh13 results in a fully penetrant XY sex-reversal phenotype, indistinguishable from that caused by complete loss of Enh13 (Gonen et al. 2018) or loss of the *Sox9* gene itself (Chaboissier et al. 2004; Lavery et al. 2011). This demonstrates that Enh13 is strictly required during the sex-determining time window to initiate the male developmental program. These findings reinforce a threshold-based model of sex determination in which Enh13 acts as the main ignition point for the elevated *Sox9* testicular expression. By integrating transient SRY input with SF1 and early chromatin accessibility, Enh13 drives *Sox9* expression above the critical level required to commit supporting cell precursors to the Sertoli cell fate. Failure to reach this threshold, as observed upon early enhancer deletion, leads to collapse of the testicular pathway, and activation of ovarian differentiation programs instead.

A central question addressed in this study was whether Enh13 continues to contribute to *Sox9* regulation after the Sertoli cell fate has been established. Late conditional gonadal deletion of Enh13 using *Amh*-Cre, which becomes active after testis cords have formed and *Sox9* expression is already robust, reveals no detectable effects on gonadal morphology, Sertoli cell identity, or adult testis maintenance. *Sox9* transcript levels remains unchanged, and mutant males are phenotypically normal and fertile. These findings indicate that Enh13 is not required for the maintenance of *Sox9* expression once the testicular program has been initiated. This stands in contrast to late conditional deletion of *Sox9* itself, which leads to progressive testicular degeneration and loss of male fertility (Barrionuevo et al. 2009), and highlights a key distinction between gene function and enhancer function. While *Sox9* remains continuously required, the enhancer responsible for its initial activation can be dispensable at later stages. This suggests that maintenance of *Sox9* expression is supported by a distinct set of regulatory elements that gain accessibility after sex determination. Indeed, chromatin accessibility analysis presented here identify multiple candidate enhancers that open in gonads at later embryonic stages and adulthood and may collectively sustain *Sox9* transcription in differentiated Sertoli cells. In human gonads, it was suggested that TESCO and the human nearby enhancer, eALDI, may function in *SOX9* maintenance rather than gene expression initiation (Croft et al. 2018). As no phenotype appeared upon late loss of Enh13 in the testis, there was no need to cross these mice to a *Sox8* null genetic background as was done in the late cKO of the *Sox9* gene itself (Barrionuevo et al. 2009).

Enhancers are generally considered to function within redundant regulatory networks, where multiple elements contribute cooperatively to ensure robust gene expression across developmental time and environmental variability (Long et al. 2016; Chatterjee and Ahituv 2017). The spatiotemporal function of enhancers is mostly characterised using accessibility techniques as ATAC-seq, combined with specific histone marks, and 3D genomic organisation methods that alter between different tissues and developmental time points (Long et al. 2016; Spitz 2016; Chatterjee and Ahituv 2017). Yet, numerous studies demonstrated that deletion of individual enhancers often produces mild or undetectable phenotypes due to compensation by so-called “shadow enhancers” or partially redundant regulatory elements (Spitz 2016; Osterwalder et al. 2018). Hence, Enh13 serves as an optimal "model enhancer" that presents both evolutionary conservation, along with a strong and clear phenotype upon its deletion (Croft et al. 2018; Gonen et al. 2018). This paves the way of using Enh13 in order to explore questions not possible to be raised in relation to other enhancers as: "Does an enhancer that is extremely potent at a given time point and tissue, continue to function later on, in the same tissue?" To the best of our knowledge, this study represents the first *in vivo* conditional knockout of a developmental enhancer, allowing enhancer function to be interrogated in a temporally controlled manner rather than through constitutive deletion at the zygote stage. Such temporal specificity would not be apparent in conventional enhancer knockout models and underscores the importance of conditional strategies for dissecting regulatory logic. This enhancer handoff model, in which early, potent enhancer establish lineage commitment and later enhancers maintain expression, is consistent with emerging principles of developmental gene regulation and provides a mechanistic framework for understanding how robustness is achieved over time (Long et al. 2016).

Human genetic data strongly implicate Enh13/eSR-A copy number variation in 46, XX testicular DSD, raising the possibility that increased Enh13 dosage alone might be sufficient to elevate *SOX9* expression above the male-determining threshold (Croft et al. 2018; Sajan et al. 2023). One possible hypothesis is that the basal levels of *Sox9*, present at the bipotential stage in XX and XY gonads (Morais da Silva et al. 1996; Zhao et al. 2018), may mediate this. However, although initially expressed in both sexes, SOX9 is actively excluded from the nucleus at that stage through an exportin-dependent nuclear export mechanism (Gasca et al. 2002). Our study shows that duplication of Enh13 in mice does not increase *Sox9* expression or induce testicular development in XX mice, nor does it alter gonadal morphology in XY mice at embryonic or adult stages. The fact that XY homozygous mice carrying the duplication present normal testis and male development suggest that even upon the duplication, Enh13 could properly function to induce *Sox9* expression or else an XY sex reversal would have been observed. The lack of XX sex reversal in mice could suggest a difference in sensitivity to enhancer dosage between mice and human. Indeed, differences in sensitivity have been reposted before where in humans, a heterozygous variant in the *SOX9* gene itself leads to Campomelic Dysplasia and 46, XY DSD (Foster et al. 1994; Wagner et al. 1994) whereas in mice, a parallel sex reversal phenotype is only seen in a homozygous state (Bi et al. 2001; Chaboissier et al. 2004; Lavery et al. 2011). This was further demonstrated when a ∼25% of normal *Sox9* levels in mice were shown as the comparable to the 50% loss of *SOX9* expression needed in human to prevent proper testis development (Gonen et al. 2017). Yet, in here, Enh13 duplication in mice does not result in even slight elevation in *Sox9* expression levels that could indicate a differences in threshold levels. Instead, it may suggest that the duplication in humans have altered spatial organization of the regulatory landscape, including three-dimensional chromatin architecture, that elevated *SOX9* in XX or prevented its repression, leading to testis development. The smallest duplication reported in humans was a 3.7 kb region (Sajan et al. 2023). It is possible that this region contained other regulatory elements, or that the duplication altered the 3D genome organisation is a way that modified *SOX9* gene expression. Interestingly, the mechanism enabling *Sox9* testicular gene expression in the SRY-deficient Amami spiny rat was shown to be mediated by male-specific duplication of yet another *Sox9* enhancer, termed Enh14 (Terao et al. 2022). Embryonic mice XX gonads in which the mouse Enh14 was replaced with the duplicated spiny rat Enh14 showed increased *Sox9* expression along with decreased *Foxl2* expression (Terao et al. 2022). Although Enh14, located ∼130 kb downstream to Enh13, has been shown to be redundant for mouse testicular *Sox9* expression (Gonen et al. 2018), the spiny rat model demonstras how duplication of an enhancer can lead to elevated gene expression, not like the current state with Enh13 duplication in mice.

Altogether, this research identifies Enh13 as an exceptional enhancer that defies several general principles of enhancer biology. Enh13 is uniquely indispensable during a critical developmental transition, acting as a gatekeeper for *Sox9* activation and testis determination. Its requirement is sharply restricted to early gonadal development, after which regulatory control is transferred to other enhancers within the *Sox9* regulatory landscape. It also highlights potential differences in enhancer transcriptional output between difference species upon copy number variation.

## Materials and Methods

### Animal Ethics Statement

All animals were maintained with appropriate husbandry according to Bar-Ilan University ethics protocols 64-11-2020, 61-11-2020, 01-01-2021, 2306-117-1 and 2305-111-1. All mice strains were maintained on a C57BL/6J genetic background. Primers used for genotyping are listed in Table 1. Frozen sperm of *Sf1*-Cre (Bingham et al. 2006) and *Amh*-Cre (Lecureuil et al. 2002) mice were kindly provided by Dr. Robin Lovell-Badge from the Francis Crick Institute. Both lines were re-derived at Bar-Ilan University transgenic unit and crossed to the newly generated Enh13 Flox allele.

**Table 1.**
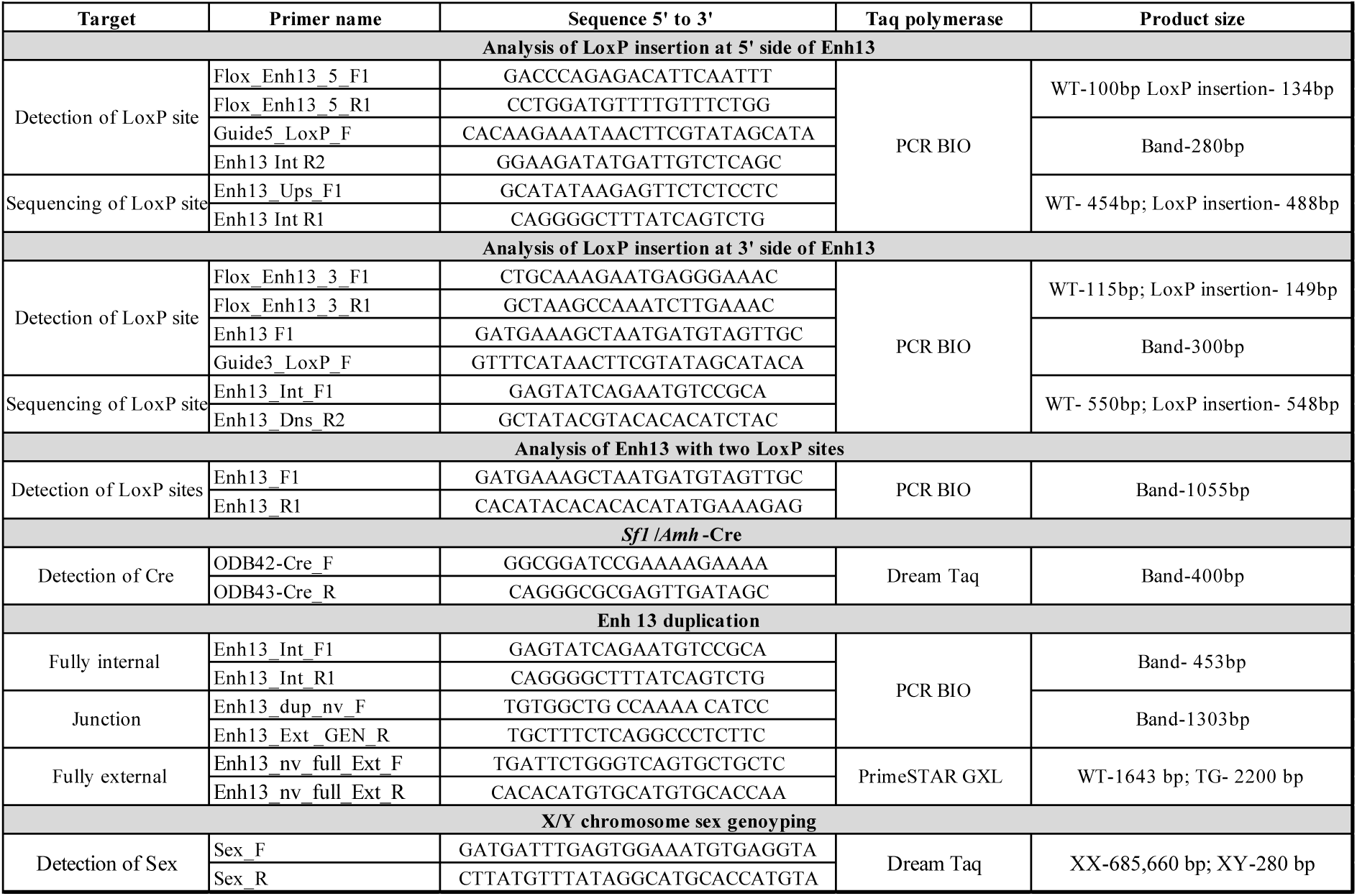
Genotyping primers.

### Design of sgRNAs mRNA and Homology Dependent Repair (HDR) single strand oligo (ssOligo)

Single guide RNAs (sgRNAs) were designed using the IDT CRISPR/Cas9 design tool (https://eu.idtdna.com). sgRNAs were chosen based on their proximity to Enh13 with the estimation that the Cas9 cuts 3-4 bp upstream of the PAM site (sgRNAs sequences are listed in Table 2). For creating crRNA-tracrRNA duplex, crRNA of all guides (IDT cat. 265065892, 265065893) and tracrRNA (IDT cat. 224893246) were resuspended into final concertation of 200µM using Nuclease Free Duplex Buffer (IDT, 11-01-03-01). crRNA and tracrRNA were mixed in 1:1 ratio, to a final complex concentration of 100µM. To anneal the sgRNA, the reaction was incubated at 95°C for 5 min, cooled to room temperature, and stored at -20°C for up to 2 weeks or until used.

**Table 2.**
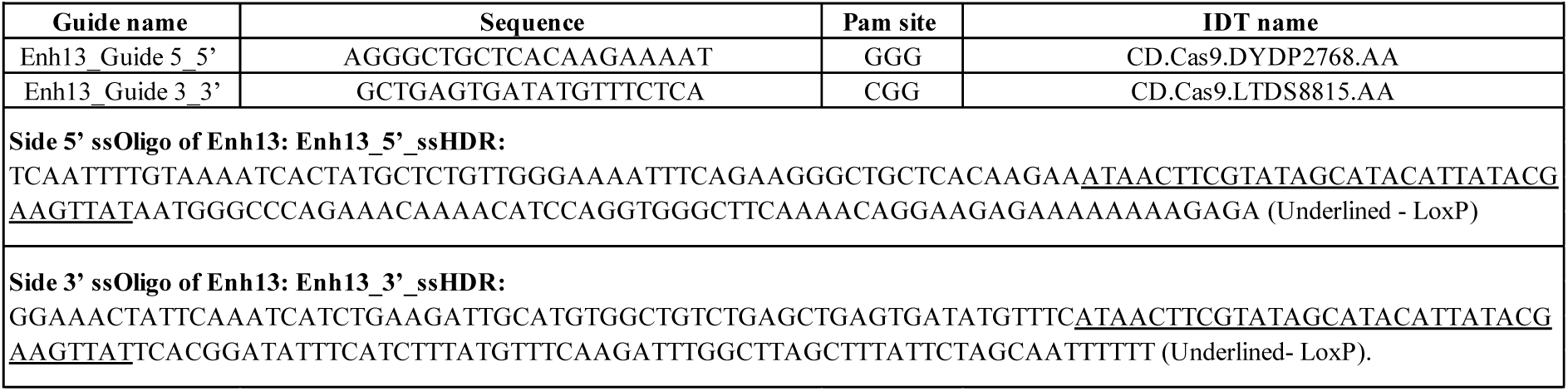
CRISPR guides and HDR sequences.

154 bp HDR donor ssOligos were ordered from IDT site and diluted in RNase free water into final concertation of 100µM (Table 2). The Oligos contain the 34 bp sequence of the LoxP and 60 bp homology arms on either side.

CRISPR/Cas9 ribonucleoproteins (RNP) assembly was performed just prior to zygote electroporation. RNP Mix was prepared within Opti-MEM (Thermo Fisher Scientific, 31985-062) and contained 1.2µM Cas9 Nuclease V3 (IDT, 1081059), 6µM sgRNA and 8 µM ssOligo. RNP Mix was incubated at room temperature for 10 minutes and then placed on ice until used for zygote electroporation.

### Zygote harvesting and CRISPR electroporation

For zygote harvesting, 4-7 weeks old C57BL6/J donor female mice were super-ovulated by administration of 5 IU of PMSG (Pregnant Mare Serum Gonadotropin; ProSpec, hor-272) using intraperitoneal (i.p.) injection. 48-50 hours following PMSG injection, females were injected with 5 IU of hCG (Human chorionic gonadotropin; Sigma Aldrich CG10). Subsequently, super-ovulated females were mated with C57BL6/J adult males in 1:1 ratio and checked for the presence of vaginal plug (VP) the morning after mating. Female mice displaying VP were sacrificed via cervical dislocation for oviduct dissection and zygote isolation. Oviducts were dissected, and ampulla nicked to release zygotes associated with surrounding cumulus cells into a M2 medium (Sigma Aldrich, M7167) containing 300 µg/ml hyaluronidase (Sigma Aldrich, H4272). Zygotes were picked using a mouth pipette and transferred to a plate containing fresh 2ml M2 and subsequently passed through several M2 washes to remove cumulus cells. Next, zygotes were moved to a plate with KSOM medium (Mercury, MR-106-D). Zygotes were kept in KSOM medium in a flat-bed CO2 incubator (5.3% CO2, 5% O2 at 37 °C) (Esco, Miri 2070047) for 30 minutes. Zygotes with the presence of polar bodies and pronucleus were isolated and washed in M2 drops followed by three washes in Opti-MEM drops and then moved to the electrode chamber containing the RNP mix. Electroporation was performed using NEPA21 electroporator (NepaGene). A large glass plate (CUY505P5 electrode with 5 mm gap) was used with 50 µl of RNP complex and 20-150 embryos. Electroporation parameters were as such: 225V poring pulse 1ms pulse width, 50 ms pulse interval and 4 repeated pulses. We then applied 20V transfer pulse, 50 ms pulse width, 50 ms pulse interval for 5 consecutive pulses. Impedance was measured before and after embryo addition as stated by the manufacturer instructions. Following electroporation, zygotes were washed with M2 and KSOM drops and surgically transferred into oviducts of pseudo-pregnant CD1 recipient females on the same day.

### In Vitro Fertilization

Sperm was harvested by nicking the apex of one cauda epididymis and transferring the sperm into a 90 μl drop of sperm capacitation media (Toyoda, Yokoyama and Hosi medium supplemented with methyl-beta-cyclodextrin (TYH+MBCD)), which was made in house according to the INFRAFRONTIER/ EMMA cryopreservation method [https://www.infrafrontier.eu/emma/cryopreservation-protocols/]. The sperm solution was incubated for 30 minutes at 37°C. In the meantime, oocytes from wild type 4-5 weeks old C57BL/6J superovulated females were collected and incubated in 200 μl drops of fertilization medium (0.25 mM glutathione in Human Tubal Fluid medium - HTF) for maximum 30 minutes. HTF media was made in house according to the INFRAFRONTIER/ EMMA protocol [https://www.infrafrontier.eu/emma/cryopreservation-protocols/]. 3-5 μl of fresh sperm was added into the fertilization medium drops containing oocytes. Three to four hours later, oocytes were washed in 4 drops of HTF, and the presumptive zygotes were used for CRISPR electroporation as described above around 9-10 hours following fertilization when polar body was evident. The following day, 2-cell stage embryos were transferred to fresh KSOM medium, and fertilization percentage was assessed. Finally, embryos were surgically transferred into oviducts of pseudo-pregnant CD1 recipient females.

### CRISPR-READI- AAV6-mediated HDR and zygote electroporation for generating Enh13 Duplication

A donor construct for HDR was generated using a custom AAV6 vector designed to introduce a tandem duplication of Enh13. The insert consisted of two consecutive 557 bp Enh13 sequences without an intervening spacer, flanked by 400 bp 5′ and 3′ homology arms (HAs), yielding a total size of 1914 bp. The sequence was synthesized and plasmid cloned using VectorBuilder and subsequently produced pilot-scale AAV6 viral stocks (Catalog #: AAV6S(VB231120-1547gue)-K1) with titter of 9.39×10^11^ GC/ml. Viral aliquots were stored at -80°C until use and thawed on ice immediately before preparation. For embryo pre-incubation with viruses, AAV6 was diluted in KSOM medium to a final concentration of 1×10^10^GC/ml. Zygotes were cultured for 6 hours with the AAVs, after which they were subjected to electroporation with RNP. Two sgRNAs targeting the endogenous 5′ and 3′ boundaries of Enh13 were used to remove the original Enh13 sequence and allow entry of the duplicated sequence instead (Table 2). RNP complex containing the two sgRNAs and Cas9 was prepared as described above and AAV pre-incubated embryos were subjected to electroporation using the NEPA21 electroporator according to the CRISPR-READI protocol (Chen et al. 2019). Electroporated embryos were surgically transferred on the same day to oviducts of pseudo-pregnant CD1 recipient females.

### Genomic DNA isolation and genotyping of genetically modified founder, F1 and F2 mice

Genomic DNA (gDNA) was extracted from tail tissue of embryos or ear punch tissue of adult animals. For embryo samples, the PCRBIO rapid extract lysis kit was used (PCRBIO, PB15.11-S). gDNA isolation from adult earpiece included 15 min incubation at 95°C with lysis buffer composed of 10mM NaOH, 0.1mM EDTA pH 8 followed by the addition of 40mM Tris HCl pH 5.

Founder mice carrying the Enh13 floxed allele were initially identified by short PCR reaction of 134 bp (5′ LoxP side) and 149 bp (3′ LoxP side) fragments allowing to identify the inserted LoxP sites, followed by Sanger sequencing of a 1055 bp fragment spanning the entire Enh13 locus containing the two LoxP sites (Table 1 for primer sequences).

Founder mice carrying the Enh13 duplication allele were first screened by PCR reactions of 453 bp fully internal PCR reaction within the insert or a 1303 bp junction PCR reaction where one primer is internal in the insert and the second is located outside the homology arm (Table 1). Next, a 2220 bp fragment was amplified encompassing the entire duplicated region and homology arms. TA cloning was performed using the pGEM®-T vector (Promega, A3600). Ligated plasmids were transformed into competent *E. coli* (NEB, C3019), plated on LB agar plates supplemented with 100 µg/mL ampicillin (Bioprep, A-221), and individual colonies were expanded for plasmid isolation using the NucleoSpin Plasmid EasyPure Kit (Macherey-Nagel, 740727). Finally, PrimeSTAR GXL DNA Polymerase (TaKaRa, R050A) was used to allow amplification across the full insertion used for Sanger sequencing (Table 1).

For Sanger sequencing, PCR products were purified using universal DNA purification kit (TIANGENE, TI-DP214-03) or by EPPiC Fast Kit (A&A biotechnologies, 1021-500F) according to the manufacturer’s instructions. Mutation analysis on the founder mice and F1 pups was done using the online DECODR (https://decodr.org/) and BLAST (https://blast.ncbi.nlm.nih.gov) tools.

Once correctly targeted founders were identified, they were crossed with C57BL/6J mice to enable germline transmission. F1 offspring were validated by Sanger sequencing, and allele-specific PCR assays were subsequently designed for routine genotyping. Heterozygous F1 males and females were bred among themselves to establish stable mutant colonies containing homozygous.

Genotyping of Enh13 *flox*, *Amh*-Cre, *Sf1*-Cre, and chromosomal sex was performed by PCR using DreamTaq 2× PCR Master Mix (Thermo Fisher Scientific, K1082) or 2× Red PCR Master Mix (PCR-BIO, PB10.23-10), according to the manufacturer’s instructions. All primer sequences are listed in Table 1.

### Time mating, gonad harvesting and gonad preparation

Embryos were collected after time mating at embryonic day E13.5 where day 0.5 was determined by the presence of vaginal plug (VP). All bright field images of gonads were taken using the Nikon Eclipse Ts2R microscope with an exposure time of 10 ms and analysed using the NIS-Elements D software.

For immunostaining, gonads from embryos, 6 weeks and 6 month postnatal mice were harvested and fixed overnight in 4% paraformaldehyde (Sigma Aldrich, P6148) in phosphate-buffered saline (PBS) at 4°C, washed three times with PBST (PBS with 0.1% Triton (Sigma Aldrich, 9002-93-1) at room temperature, incubated with 20% sucrose (Fisher BioReagents, BP220-1) overnight at 4°C and then embedded in OCT (Leica, 14020108926) and stored at - 80°C until further use.

### Immunofluorescence staining

Immunofluorescence staining of E13.5, 6 weeks and 6 months gonads were performed on 10μm-thick sagittal cryostat sections (Leica, CM3050-S). Antigen retrieval was performed for embryonic samples with DAKO (Target retrieval solution, Agilent, S1699) at 65°C for 30 min. Samples were then blocked in PBST containing 10% donkey serum (Sigma Aldrich, D9663) for 1 hour and incubated with primary antibodies (diluted in PBST containing 1% donkey serum) overnight at 4°C (All primary and secondary antibodies as well as dyes used are listed in Table 3). Following three PBST washes, secondary antibodies were added for 1 hour at room temperature (RT). Slides were then washed, dried, and mounted (Polysciences 18606-20). All immunofluorescence slides were also stained with 4’,6-diamidino-2-phenylindole (DAPI; Invitrogen, D1306) to visualize nuclear DNA. Imacges were obtained with a Leica Microsystems SP8 confocal microscope.

**Table 3.** Antibodies/ Dyes used in this study.

| Table 3. Antibodies/ Dyes used in this study |  |  |  |  |  |  |  |
| --- | --- | --- | --- | --- | --- | --- | --- |
| Target | Host | Manufacture | Cat. number | Dilution | Antibody type | Used for staining | Cell type |
| αSOX9 | Rabbit | Millipore | AB5535 | 1:2000 | Primary | Adult gonads (6W;6M) | Sertoli cells |
| αSF1 | Rat | Cosmo Bio | KAL-KO610 | 1:300 | Primary | Adult gonads (6W;6M) | Sertoli and Leydig cells |
| αDDX4 | Mouse | Abcam | ab27591 | 1:300 for males /<br>1:2000 for females | Primary | Adult gonads (6W;6M) | Germ cells |
| αSOX9 | Goat | R&D systems | AF3075 | 1:300 | Primary | Adult and Embryonic gonads (6W;E13.5) | Sertoli cells |
| αFOXL2 | Rabbit | Abcam | ab246511 | 1:300 | Primary | Adult and Embryonic gonads (6W;E13.5) | Granulosa cells |
| TNP-1 | Rabbit | Abcam | ab73135 | 1:1000 | Primary | Adult gonads (6W;6M) | Elongated spermatids |
| BOULE | Goat | LSBio | LS-B7334-50 | 1:100 | Primary | Adult gonads (6W;6M) | Spermatocytes |
| αGCNA1 (TRA98) | Rat | Abcam | ab82527 | 1:200 | Primary | Adult and Embryonic gonads (6W;E13.5) | Germ cells |
| αRat-Alexa flour488 | Donkey | Abcam | AB150153 | 1:500 | Secondary | Adult gonads (6W;6M) | - |
| αMouse-Alexa flour 647 | Donkey | Invitrogen | ab150111 | 1:500 | Secondary | Adult gonads (6W;6M) | - |
| αGoat-Alexa flour 488 | Donkey | Invitrogen | A-11055 | 1:500 | Secondary | Adult and Embryonic gonads (6W;E13.5) | - |
| αRabbit-Alexa flour 568 | Donkey | Invitrogen | A-10042 | 1:500 | Secondary | Adult and Embryonic gonads (6W;6M;E13.5) | - |
| αRat-Alexa flour 647 | Donkey | Abcam | ab150155 | 1:500 | Secondary | Adult and Embryonic gonads (6W;E13.5) | - |
| DAPI | - | Invitrogen | D-1306 | 300nM | Secondary | All | Nucleic acid |

### H&E staining of cryosections

For H&E staining, OCT cryosection slides were thawed and fixed in Acetone (Macron, C6776-25) at -20°C for 3 minutes and 80% Methanol (Merck 67-56-1) at 4°C for 5 minutes followed by PBS wash for 10 minutes. Filtered Harris Hematoxylin (Kaltek, 1513) was used for 1 minute staining followed by tap water wash for 5 minutes and then Eosin (Sigma-Aldrich, HT110332) was used for 30 seconds staining. De-hydration was performed by washing with 70%, 80%, 90% and 100% Ethanol (Macron C677716), each wash for 5 minutes. Final de-hydration and polishing was performed using 2 Xylene (Sigma-Aldrich, 534056) washes, each for 5 minutes. Slides were dried, wet again with Xylene, mounted with drops of Permount mounting medium (Fisher Chemical, SP15-100) and covered. H&E images were taken using Nikon TS100.

### RNA isolation, cDNA preparation, and Quantitative Real-Time Polymerase Chain Reaction (qRT-PCR)

Total RNA was extracted from 6w gonads using EURX GeneMATRIX Universal RNA Purification Kit (EURX, E3598) and from E13.5 gonad pairs using Qiagen RNeasy Plus micro kit (Qiagen, 74034). RNA yield was quantified with a NanoDrop spectrophotometer. For adult samples, 1µg of RNA was taken from 6 weeks old gonads for cDNA preparation using the qScript cDNA Synthesis Kit (QuantaBio 95047-100-2) according to the manufacturer’s instructions. qRT-PCR reactions were performed in duplicate using qPCR-PerfeCTa SYBR Green FastMix (95074-012-2, QuantaBio) with 140 nM each of forward and reverse primers (Table 4) and analyzed on the QuantStudio 1 Real-Time PCR System (Thermo Scientific). For embryonic samples, 200ng of RNA was taken for cDNA preparation using SuperScript™ III Reverse Transcriptase (Thermo Scientific, 18080085) according to the manufacturer’s instructions. qRT-PCR reactions were performed in duplicate using PowerSYBR Green PCR master mix (Thermo Scientific, AB-4367659) with 140 nM each of forward and reverse primers (Table 4) and analyzed on the QuantStudio 1 Real-Time PCR System (Thermo Scientific). Analysis was done using Comparative CT (2^-ΔΔCT^) technique and expression is relative to the house keeping gene, *Hprt.* Statistical analyses were carried out using Prism 10 software (GraphPad) using one-way ANOVA test followed by Dunnett’s post-test. P value < 0.05 was considered as statistically significant. The number of gonads analyzed at each stage is depicted in the graphs.

**Table 4.**
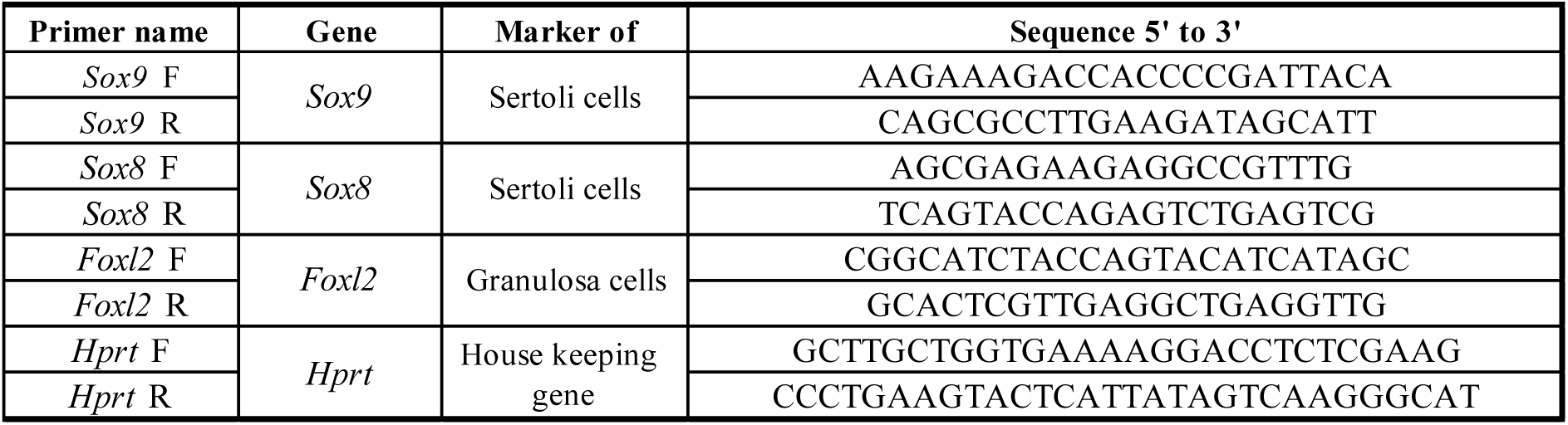
Primers used for quantitative RT-PCR.

### ATAC-seq analysis

Fastq files for ATAC-seq data from purified E10.5 gonadal somatic cells (*Nr5a1*-GFP+ cells) were obtained from (Garcia-Moreno et al. 2019) (GSE118755). ATAC-seq data from purified granulosa (*Enh8*-mCherry+) and Sertoli (*Sox9*-IRES-GFP+) cells from E11.5 to E15.5 were obtained from (Stevant et al. 2025) (GSE277646). Finally, ATAC-seq from purified postnatal granulosa cells (P23, manual follicle dissociation), and Sertoli cells (P7, *Dhh-Cre; CAG-Stop^flox^-tdTomato*) were obtained from (Lindeman et al. 2021) (GSE154484).

The downloaded FastQ files were processed together using nf-core/atacseq pipeline v2.1.2, as described in (Stevant et al. 2025). Briefly, read quality controls were performed with FastQC. Sequencing adaptors were removed with Cutadapt. Reads were mapped on the mm10/GRCm38 reference genome from Gencode (M25) with BWA. Mapped reads were filtered to remove the unpaired reads, the mitochondrial reads, the duplicated reads, the multimapped reads, the fragments with insert size > 2 kb, and the reads mapping to the blacklisted regions (https://github.com/Boyle-Lab/Blacklist/blob/master/lists/mm10-blacklist.v2.bed.gz).

Peaks were called with MACS2 using the “narrow_peaks” parameter. For each condition, peaks with FDR<0.01 and found in at least two replicates were merged as consensus open chromatin regions. The obtained consensus regions were then combined and merged to constitute the set of non-overlapping open chromatin regions present in any conditions and was used for the downstream analysis. Bigwig files normalized by million mapped reads from the nf-core/atacseq pipeline were corrected to be normalized by the size factors (Variance Stabilizing Transformation) calculated from reads in peaks using DESeq2 (https://doi.org/10.1186/s13059-014-0550-8) in order to homogenize the signal from the different datasets.

*Sox9* enhancer annotation was taken from (Gonen et al. 2018) and converted from mm9 to mm10 using LiftOver (UCSC website). ATAC signal was visualized using the Gviz R package (https://doi.org/10.1007/978-1-4939-3578-9_16).

## Competing declaration

The authors declare that they have no competing interests.

## Acknowledgements

We thank the BIU animal facility and technicians for the help with animal maintenance. We are grateful to the BIU transgenic facility for production of CRISPR genome-edited mice. We acknowledge the Life Sciences Microscopy unit at BIU for help with imaging.

## Author contribution

ML, MR and NG conceived the idea and experiments; ML, MR, YCD and EA performed experiments and analysed the data. IS re-analysed the ATAC-seq data. SZL and ML generated the CRISPR-genome edited mice. The manuscript was written by NG, ML, MR and EA. All authors read and accepted the data being presented in the manuscript.

## Data and materials availability

All data is available in the manuscript or the supplementary materials.

---

**Supplemental Figure 1.**
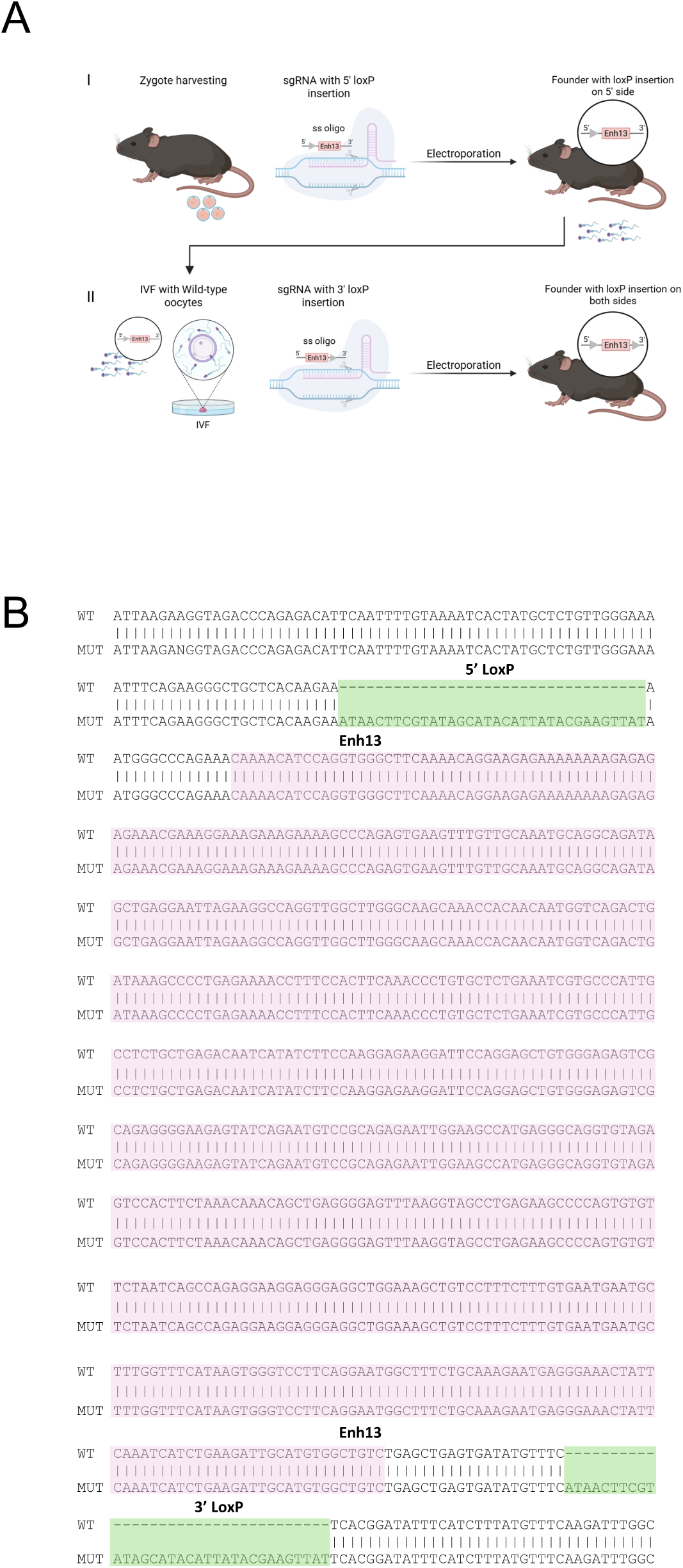
Targeting approach and validation of the Enh13 floxed allele. (A) Schematic representation of the targeting approach for generating a floxed allele of Enh13. In phase Ι, LoxP insertion on the 5’ side of Enh13 is carried via electroporation on WT zygotes resulting in founder mice with LoxP inserted on the 5’ side of Enh13. In phase ΙΙ, sperm isolation was performed from a germline transmitted mouse, homozygous for 5’ LoxP insertion, and zygote were generated using IVF. LoxP was inserted on the 3’ side by electroporation generating mice with LoxP insertion on both the 5’ and 3’ sides of Enh13 (B) Representative BLAST alignment between WT Enh13 sequence and homozygous mouse carrying the 5′ and 3′ LoxP sequence outside Enh13. WT sequences are shown on the top lines and mutant (MUT) sequences on the bottom lines. LoxP sequences are highlighted in green, and Enh13 in pink.

**Supplemental Figure 2.**
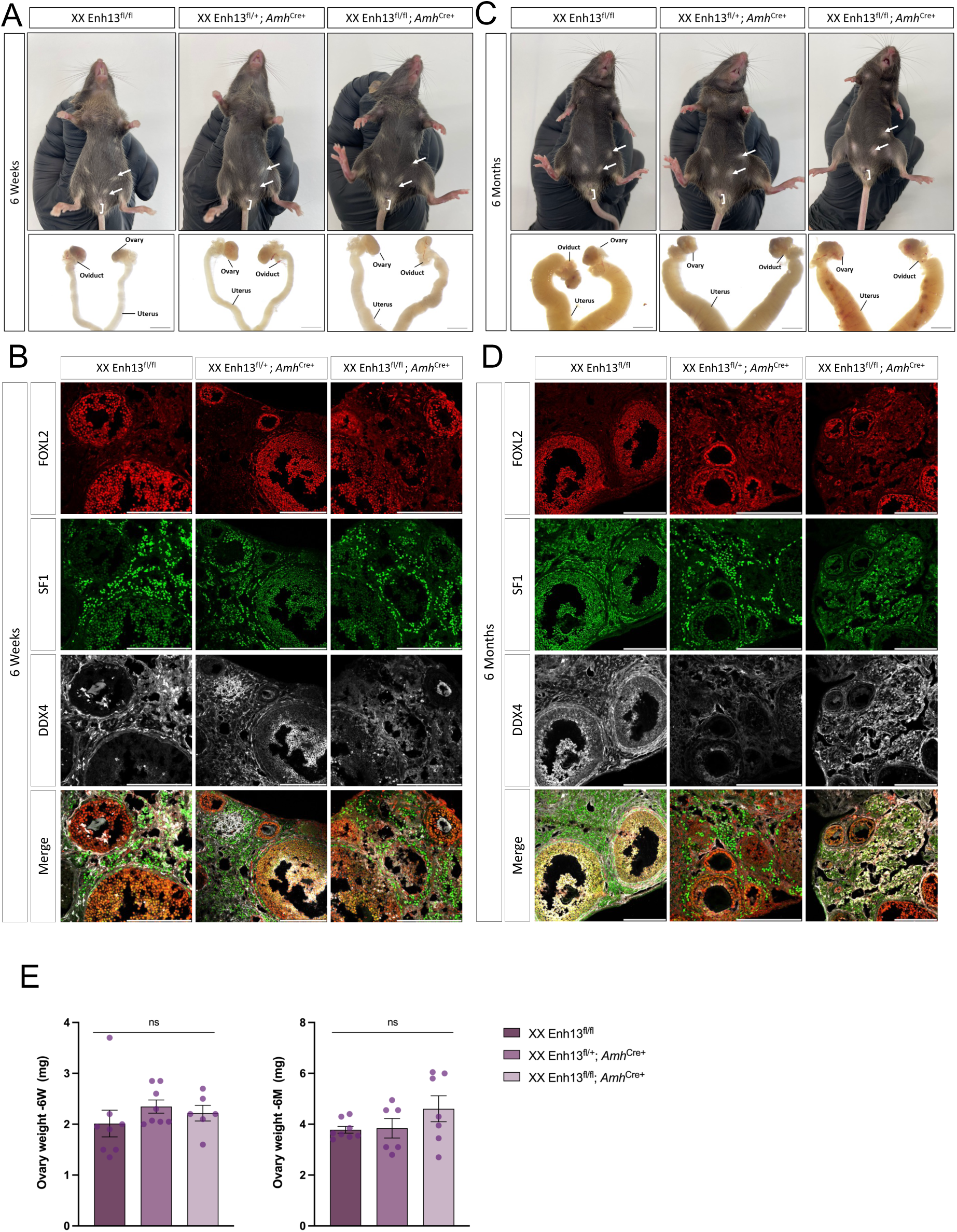
Characterization of Enh13^fl/fl^; *Amh*^Cre^ mouse line in females. (A, C) External genitalia and brightfield images of internal reproductive organs of XX Enh13 ^fl/fl^, XX Enh13 ^fl/+^; *Amh*^Cre+^, and XX Enh13 ^fl/fl^; *Amh*^Cre+^ mice at 6-Week-old (A) and at 6-Months-old (C). Scale bars represent 2000 μm. (B, E) Immunostaining of gonads from 6-Week-old (B) and 6-Month-old mice (E). Gonads were stained for granulosa-marker FOXL2 (red), theca cell marker SF1 (green) and germ cells marker DDX4 (grey). Scale bars represent 200 μm. (D) Quantification of ovary weight from 6-Week-old mice and 6-Mmonth-old mice. Each point represents the average weight of both ovaries of an individual mouse; error bars show SEM. Statistical analysis was performed using one-way ANOVA tests in Prism 10 software. P values below 0.05 were considered statistically significant. ns- not significant.

**Supplemental Figure 3.**
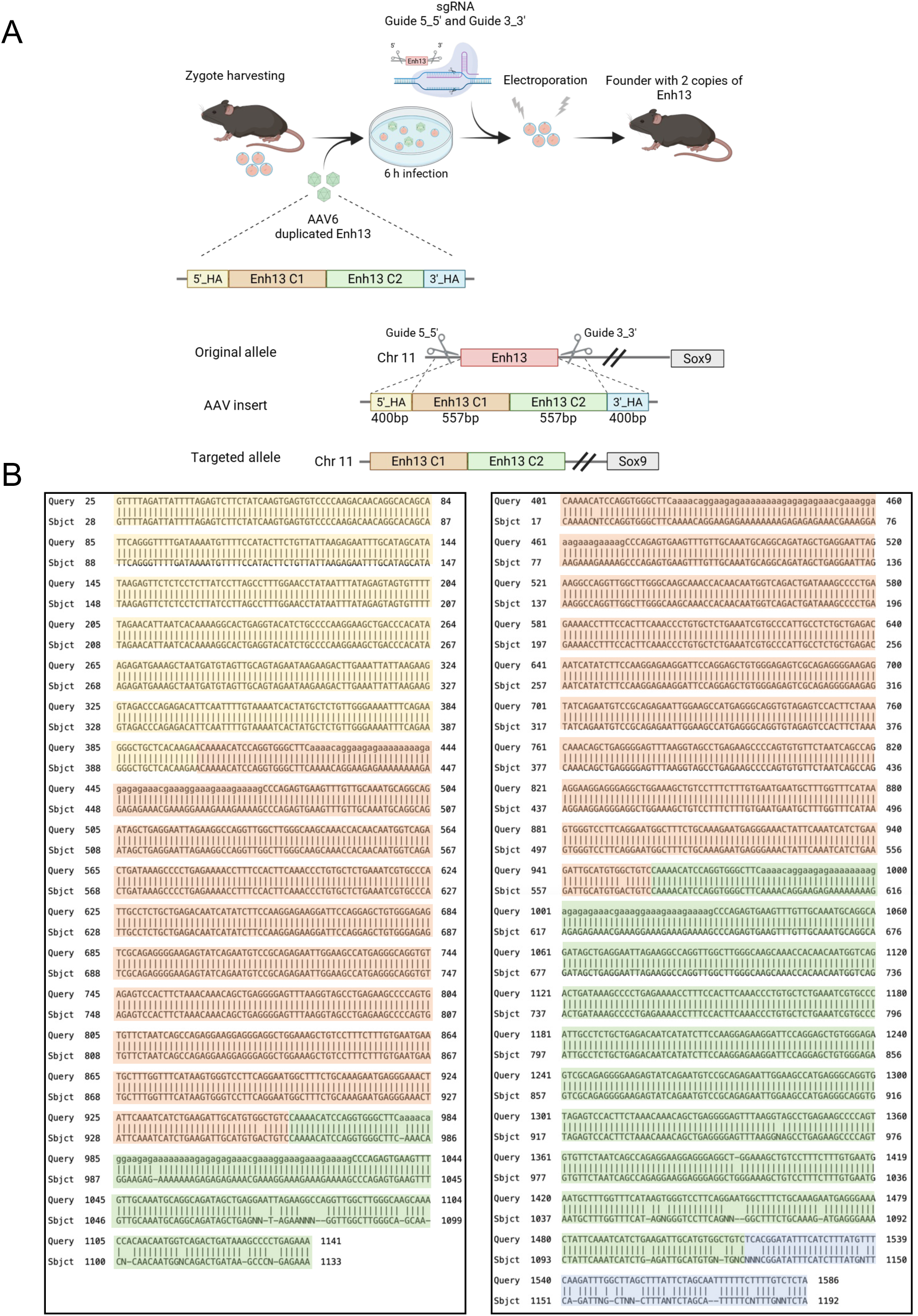
Targeting approach and validation of the Enh13 duplicated allele. (A) Schematic representation of the targeting approach using the CRISPR-READI workflow. Zygotes are harvested from super ovulated female mice, pre-incubated with rAAV6 harboring the donor template containing two copies of Enh13 and homology arms, electroporated with Cas9/sgRNA RNPs to delete the original Enh13, and implanted into pseudo pregnant females to generate founder edited mice. Schematic representation of the cleavage sites to remove the original Enh13 locus and below is the insert of duplicated Enh13 flanked by 400 bp homology arms. The targeted allele containing two copies of Enh13 instead of the original one is presented below. (B) Sanger sequencing alignment of Enh13 duplication insertion sites analyzed using the BLAST tool. Predicted duplicated sequences (Query) are shown on the top lines and actual F2 homozygous mice sequences (Sbjct) on the bottom lines. 5’ HA sequences are highlighted in yellow, Enh13 copy 1-orange, Enh13 copy 2-green and 3’ HA in blue.

**Supplemental Figure 4.**
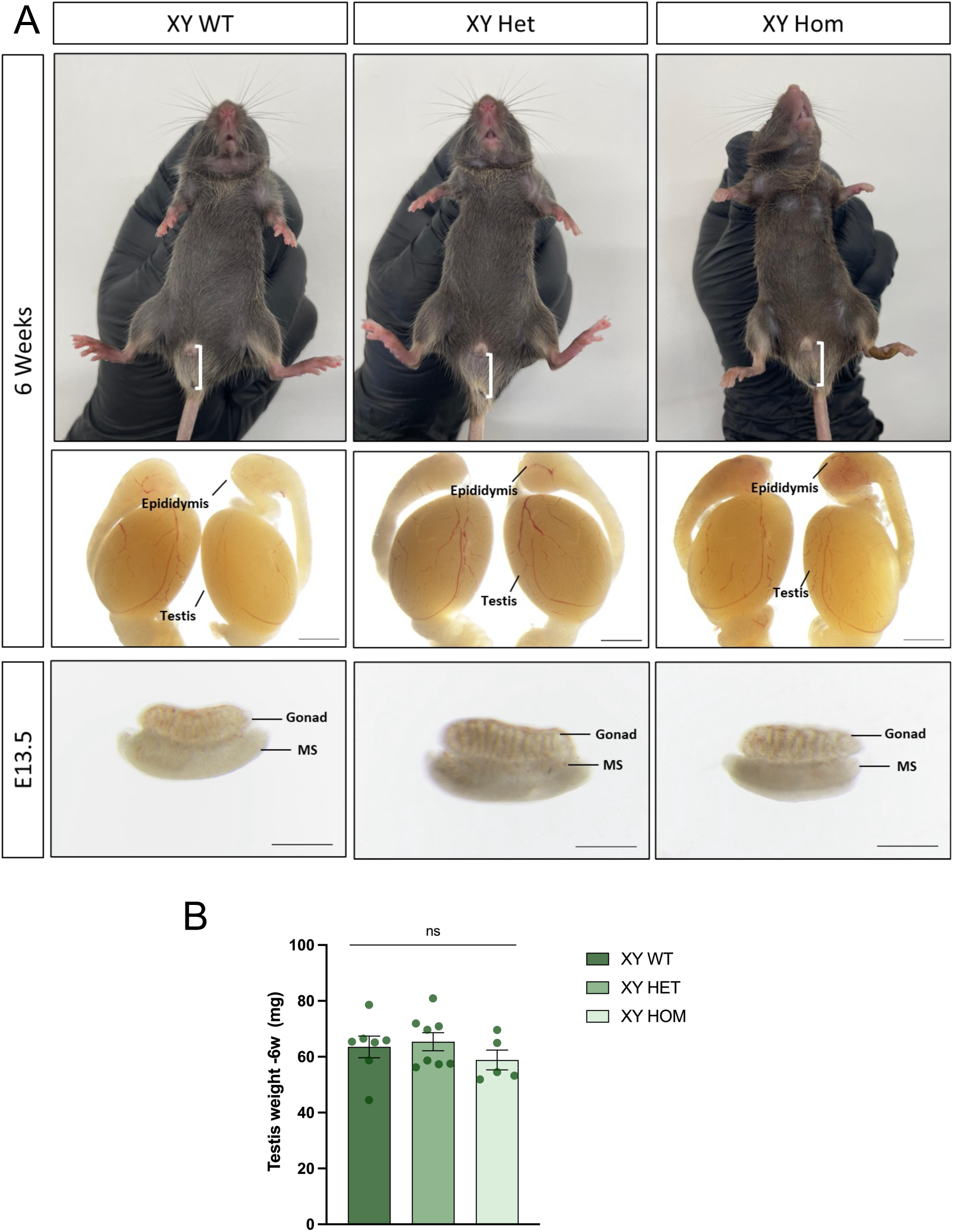
Characterization of XY mice carrying Enh13 duplication. (A) Bright field images of the external and internal genitalia and gonads of 6-Week-old adult and E13.5 embryonic gonads of XY WT, heterozygous and homozygous mice for Enh13 duplication. Scale Bar represents 2000 µm in adults and 500 µm in embryos. (B) Quantification of testis weight in 6-Week-old mice. Each dot represents average weight of both tests of an individual mouse; error bars show SEM. Statistical analysis was performed using one-way ANOVA test in Prism 10 software where samples were compares to XX WT. P values below 0.05 were considered statistically significant. ns- not significant.

## References

Baetens D, Mendonca BB, Verdin H, Cools M, De Baere E. 2017. Non-coding variation in disorders of sex development. Clin Genet 91: 163–172.

Barrionuevo F, Georg I, Scherthan H, Lecureuil C, Guillou F, Wegner M, Scherer G. 2009. Testis cord differentiation after the sex determination stage is independent of Sox9 but fails in the combined absence of Sox9 and Sox8. Dev Biol 327: 301–312.

Bi W, Huang W, Whitworth DJ, Deng JM, Zhang Z, Behringer RR, de Crombrugghe B. 2001. Haploinsufficiency of Sox9 results in defective cartilage primordia and premature skeletal mineralization. Proc Natl Acad Sci U S A 98: 6698–6703.

Bingham NC, Verma-Kurvari S, Parada LF, Parker KL. 2006. Development of a steroidogenic factor 1/Cre transgenic mouse line. Genesis 44: 419–424.

Bolt CC, Duboule D. 2020. The regulatory landscapes of developmental genes. Development 147.

Chaboissier MC, Kobayashi A, Vidal VI, Lutzkendorf S, van de Kant HJ, Wegner M, de Rooij DG, Behringer RR, Schedl A. 2004. Functional analysis of Sox8 and Sox9 during sex determination in the mouse. Development 131: 1891–1901.

Chatterjee S, Ahituv N. 2017. Gene Regulatory Elements, Major Drivers of Human Disease. Annu Rev Genomics Hum Genet 18: 45–63.

Chen S, Sun S, Moonen D, Lee C, Lee AY, Schaffer DV, He L. 2019. CRISPR-READI: Efficient Generation of Knockin Mice by CRISPR RNP Electroporation and AAV Donor Infection. Cell Rep 27: 3780–3789 e3784.

Croft B, Ohnesorg T, Hewitt J, Bowles J, Quinn A, Tan J, Corbin V, Pelosi E, van den Bergen J, Sreenivasan R et al. 2018. Human sex reversal is caused by duplication or deletion of core enhancers upstream of SOX9. Nat Commun 9: 5319.

Foster JW, Dominguez-Steglich MA, Guioli S, Kwok C, Weller PA, Stevanovic M, Weissenbach J, Mansour S, Young ID, Goodfellow PN et al. 1994. Campomelic dysplasia and autosomal sex reversal caused by mutations in an SRY-related gene. Nature 372: 525–530.

Franke M, Ibrahim DM, Andrey G, Schwarzer W, Heinrich V, Schopflin R, Kraft K, Kempfer R, Jerkovic I, Chan WL et al. 2016. Formation of new chromatin domains determines pathogenicity of genomic duplications. Nature 538: 265–269.

Garcia-Moreno SA, Futtner CR, Salamone IM, Gonen N, Lovell-Badge R, Maatouk DM. 2019. Gonadal supporting cells acquire sex-specific chromatin landscapes during mammalian sex determination. Dev Biol 446: 168–179.

Gasca S, Canizares J, De Santa Barbara P, Mejean C, Poulat F, Berta P, Boizet-Bonhoure B. 2002. A nuclear export signal within the high mobility group domain regulates the nucleocytoplasmic translocation of SOX9 during sexual determination. Proc Natl Acad Sci U S A 99: 11199–11204.

Gonen N, Futtner CR, Wood S, Garcia-Moreno SA, Salamone IM, Samson SC, Sekido R, Poulat F, Maatouk DM, Lovell-Badge R. 2018. Sex reversal following deletion of a single distal enhancer of Sox9. Science 360: 1469–1473.

Gonen N, Lovell-Badge R. 2019. The regulation of Sox9 expression in the gonad. Curr Top Dev Biol 134: 223–252.

Gonen N, Quinn A, O’Neill HC, Koopman P, Lovell-Badge R. 2017. Normal Levels of Sox9 Expression in the Developing Mouse Testis Depend on the TES/TESCO Enhancer, but This Does Not Act Alone. PLoS Genet 13: e1006520.

Huang B, Wang S, Ning Y, Lamb AN, Bartley J. 1999. Autosomal XX sex reversal caused by duplication of SOX9. Am J Med Genet 87: 349–353.

Ibrahim DM, Mundlos S. 2020. Three-dimensional chromatin in disease: What holds us together and what drives us apart? Curr Opin Cell Biol 64: 1–9.

Kim Y, Bingham N, Sekido R, Parker KL, Lovell-Badge R, Capel B. 2007. Fibroblast growth factor receptor 2 regulates proliferation and Sertoli differentiation during male sex determination. Proc Natl Acad Sci U S A 104: 16558–16563.

Kim Y, Kobayashi A, Sekido R, DiNapoli L, Brennan J, Chaboissier MC, Poulat F, Behringer RR, Lovell-Badge R, Capel B. 2006. Fgf9 and Wnt4 act as antagonistic signals to regulate mammalian sex determination. PLoS Biol 4: e187.

Koopman P, Gubbay J, Vivian N, Goodfellow P, Lovell-Badge R. 1991. Male development of chromosomally female mice transgenic for Sry. Nature 351: 117–121.

Lavery R, Lardenois A, Ranc-Jianmotamedi F, Pauper E, Gregoire EP, Vigier C, Moreilhon C, Primig M, Chaboissier MC. 2011. XY Sox9 embryonic loss-of-function mouse mutants show complete sex reversal and produce partially fertile XY oocytes. Dev Biol 354: 111–122.

Lecureuil C, Fontaine I, Crepieux P, Guillou F. 2002. Sertoli and granulosa cell-specific Cre recombinase activity in transgenic mice. Genesis 33: 114–118.

Lindeman RE, Murphy MW, Agrimson KS, Gewiss RL, Bardwell VJ, Gearhart MD, Zarkower D. 2021. The conserved sex regulator DMRT1 recruits SOX9 in sexual cell fate reprogramming. Nucleic Acids Res 49: 6144–6164.

Liu CF, Samsa WE, Zhou G, Lefebvre V. 2017. Transcriptional control of chondrocyte specification and differentiation. Semin Cell Dev Biol 62: 34–49.

Long HK, Osterwalder M, Welsh IC, Hansen K, Davies JOJ, Liu YE, Koska M, Adams AT, Aho R, Arora N et al. 2020. Loss of Extreme Long-Range Enhancers in Human Neural Crest Drives a Craniofacial Disorder. Cell Stem Cell 27: 765–783 e714.

Long HK, Prescott SL, Wysocka J. 2016. Ever-Changing Landscapes: Transcriptional Enhancers in Development and Evolution. Cell 167: 1170–1187.

Morais da Silva S, Hacker A, Harley V, Goodfellow P, Swain A, Lovell-Badge R. 1996. Sox9 expression during gonadal development implies a conserved role for the gene in testis differentiation in mammals and birds. Nat Genet 14: 62–68.

O’Bryan MK, Takada S, Kennedy CL, Scott G, Harada S, Ray MK, Dai Q, Wilhelm D, de Kretser DM, Eddy EM et al. 2008. Sox8 is a critical regulator of adult Sertoli cell function and male fertility. Dev Biol 316: 359–370.

Ogawa Y, Terao M, Hara S, Tamano M, Okayasu H, Kato T, Takada S. 2018. Mapping of a responsible region for sex reversal upstream of Sox9 by production of mice with serial deletion in a genomic locus. Sci Rep 8: 17514.

Ogawa Y, Tsuchiya I, Yanai S, Baba T, Morohashi KI, Sasaki T, Sasaki J, Terao M, Tsuji-Hosokawa A, Takada S. 2025. Author Correction: GATA4 binding to the Sox9 enhancer mXYSRa/Enh13 is critical for testis differentiation in mouse. Commun Biol 8: 199.

Osterwalder M, Barozzi I, Tissieres V, Fukuda-Yuzawa Y, Mannion BJ, Afzal SY, Lee EA, Zhu Y, Plajzer-Frick I, Pickle CS et al. 2018. Enhancer redundancy provides phenotypic robustness in mammalian development. Nature 554: 239–243.

Ridnik M, Abberbock E, Alipov V, Lhermann SZ, Kaufman S, Lubman M, Poulat F, Gonen N. 2024. Two redundant transcription factor binding sites in a single enhancer are essential for mammalian sex determination. Nucleic Acids Res 52: 5514–5528.

Rotgers E, Jorgensen A, Yao HH. 2018. At the Crossroads of Fate-Somatic Cell Lineage Specification in the Fetal Gonad. Endocr Rev 39: 739–759.

Sajan SA, Brown CM, Davis-Keppen L, Burns K, Royer E, Coleman JAC, Hilton BA, DuPont BR, Perry DL, Taft RJ et al. 2023. The smallest likely pathogenic duplication of a SOX9 enhancer identified to date in a family with 46,XX testicular differences of sex development. Am J Med Genet A.

Schmahl J, Kim Y, Colvin JS, Ornitz DM, Capel B. 2004. Fgf9 induces proliferation and nuclear localization of FGFR2 in Sertoli precursors during male sex determination. Development 131: 3627–3636.

Schoenfelder S, Fraser P. 2019. Long-range enhancer-promoter contacts in gene expression control. Nat Rev Genet 20: 437–455.

Sekido R, Bar I, Narvaez V, Penny G, Lovell-Badge R. 2004. SOX9 is up-regulated by the transient expression of SRY specifically in Sertoli cell precursors. Dev Biol 274: 271–279.

Sekido R, Lovell-Badge R. 2008. Sex determination involves synergistic action of SRY and SF1 on a specific Sox9 enhancer. Nature 453: 930–934.

Sekido R, Lovell-Badge R. 2009. Sex determination and SRY: down to a wink and a nudge? Trends Genet 25: 19–29.

Sinclair AH, Berta P, Palmer MS, Hawkins JR, Griffiths BL, Smith MJ, Foster JW, Frischauf AM, Lovell-Badge R, Goodfellow PN. 1990. A gene from the human sex-determining region encodes a protein with homology to a conserved DNA-binding motif. Nature 346: 240-244.

Sock E, Schmidt K, Hermanns-Borgmeyer I, Bosl MR, Wegner M. 2001. Idiopathic weight reduction in mice deficient in the high-mobility-group transcription factor Sox8. Mol Cell Biol 21: 6951–6959.

Spitz F. 2016. Gene regulation at a distance: From remote enhancers to 3D regulatory ensembles. Semin Cell Dev Biol 57: 57–67.

Sreenivasan R, Gonen N, Sinclair A. 2022. SOX Genes and Their Role in Disorders of Sex Development. Sex Dev 16: 80–91.

Stevant I, Abberbock E, Ridnik M, Weiss R, Swisa L, Futtner CR, Maatouk DM, Lovell-Badge R, Malysheva V, Gonen N. 2025. The gene regulatory landscape driving mouse gonadal supporting cell differentiation. Sci Adv 11: eadv1885.

Stevant I, Nef S. 2019. Genetic Control of Gonadal Sex Determination and Development. Trends Genet 35: 346–358.

Terao M, Ogawa Y, Takada S, Kajitani R, Okuno M, Mochimaru Y, Matsuoka K, Itoh T, Toyoda A, Kono T et al. 2022. Turnover of mammal sex chromosomes in the Sry-deficient Amami spiny rat is due to male-specific upregulation of Sox9. Proc Natl Acad Sci U S A 119: e2211574119.

Vidal VP, Chaboissier MC, de Rooij DG, Schedl A. 2001. Sox9 induces testis development in XX transgenic mice. Nat Genet 28: 216–217.

Wagner T, Wirth J, Meyer J, Zabel B, Held M, Zimmer J, Pasantes J, Bricarelli FD, Keutel J, Hustert E et al. 1994. Autosomal sex reversal and campomelic dysplasia are caused by mutations in and around the SRY-related gene SOX9. Cell 79: 1111–1120.

Wilhelm D, Martinson F, Bradford S, Wilson MJ, Combes AN, Beverdam A, Bowles J, Mizusaki H, Koopman P. 2005. Sertoli cell differentiation is induced both cell-autonomously and through prostaglandin signaling during mammalian sex determination. Dev Biol 287: 111–124.

Wilhelm D, Palmer S, Koopman P. 2007. Sex determination and gonadal development in mammals. Physiol Rev 87: 1–28.

Wilhelm D, Perea-Gomez A, Newton A, Chaboissier MC. 2025. Gonadal sex determination in vertebrates: rethinking established mechanisms. Development 152.

Yang Y, Gomez N, Infarinato N, Adam RC, Sribour M, Baek I, Laurin M, Fuchs E. 2023. The pioneer factor SOX9 competes for epigenetic factors to switch stem cell fates. Nat Cell Biol 25: 1185–1195.

Zhao L, Wang C, Lehman ML, He M, An J, Svingen T, Spiller CM, Ng ET, Nelson CC, Koopman P. 2018. Transcriptomic analysis of mRNA expression and alternative splicing during mouse sex determination. Mol Cell Endocrinol.

